# Chemosensory diversity across the order Vibrionales reveals a conserved core and three accessory signalling systems

**DOI:** 10.64898/2026.05.11.724241

**Authors:** Shravani Rajan Rawool, Gaurav Sharma

## Abstract

Chemosensory systems (CSSs) are multi-protein assemblies regulating bacterial motility and diverse alternative cellular functions. Despite extensive study of chemotaxis in *Vibrio cholerae*, a comprehensive evolutionary analysis of CSSs across order Vibrionales has been lacking. Using 116 curated representative genomes across 28 *Vibrio* clades and ∼10,000 RefSeq/metagenome-assembled genomes, we characterized the chemosensory toolkit of Vibrionales. CheA-based phylogenetics, CSS architecture, sequence similarity networks, structural comparisons, and synteny analysis identified four discrete CSS types: F6, F7, F9, and an experimentally unexplored F8. F6 is universally conserved on chromosome I and essential for flagellar motility (chemotaxis), while F7, F8, and F9 show patchy, replicon-flexible distributions reflecting lineage-specific retention or horizontal acquisition. F6, F7, and F8 were vertically inherited from Gammaproteobacteria and retained in selected species, whereas F9 was horizontally acquired from Alphaproteobacteria in only selected species. Structural analysis further reveals conserved CheA folds despite sequence divergence, with lineage-specific insertions in F8 and F9 CheA proteins. Collectively, this study reveals a two-tier chemosensory architecture within order Vibrionales: 1) a chromosomally stable F6 motility core under purifying selection, 2) overlaid by dynamically evolving F7, F8, and F9 accessory systems, wherein multipartite genome organization might serve as an evolutionary key for sensory innovation in order Vibrionales, enabling rapid niche adaptation without compromising core chemotactic fidelity.

**Importance:** Bacterial chemosensory systems enable cells to detect and respond to a wide range of environmental stimuli, coordinating cellular behaviours such as motility, surface attachment, biofilm formation, and other adaptive responses. CSS system helps bacteria to colonize the host environment along with their virulence mechanism, yet how these systems diversified and evolved across the broader *Vibrio* family remained unknown. By analysing over 10,000 *Vibrio* genomes, this study maps CSS diversity order-wide and reveals how the bacteria’s distinctive two-chromosome genome enables flexible, niche-tailored sensory assembly.

## Introduction

Bacteria are present omnipotently in nature, ranging from free-living to being associated with plants and animals in symbiotic/pathogenic associations. Order Vibrionales members exemplify this ecological versatility as their habitat ranges from free-living, pathogenic, symbiotic, to commensal. Along with important human pathogens such as *Vibrio cholerae*, *V. parahaemolyticus*, and *V. vulnificus*, *Aliivibrio fischeri* forms a well-known mutualistic association with marine squid, where its bioluminescence helps the host avoid predators (Miyashiro & Ruby, 2012). Other members, such as *V. anguillarum*, *V. harveyi*, and *V. coralliilyticus*, are well-known marine host-pathogens. Taxonomically, this order belongs to class Gammaproteobacteria within phylum Pseudomonadota (formerly Proteobacteria) and comprises a single family, Vibrionaceae, which includes 15 representative genera: *Allivibrio, Allocatenococcus, Allomonas*, *Photobacterium*, *Candidatus Photodemus, Corallibacterium, Thaumasivibrio, Paraphotobacterium, Veronia, Enterovibrio, Vibrio, Grimontia, Salinivibrio, Parasalinivibrio,* and *Photococcus*. Over 190 species are currently recognized within the family, with phylogenetic analyses resolving them into distinct evolutionary clades (Jiang et al., 2022). These Gram-negative, facultatively anaerobic bacteria typically possess one or more polar flagella and characteristic multipartite genomes comprising two chromosomes and occasional plasmids or megaplasmids.

The ability of Vibrionales organisms to thrive in highly variable environments (such as marine, diverse hosts, etc.) depends on sophisticated signal-transduction mechanisms that enable cells to sense and respond to nutrients. Bacterial signal transduction encompasses diverse regulatory architectures, ranging from one-component systems (OCSs) and two-component systems (TCSs) to more complex signaling networks such as quorum sensing and chemosensory systems (CSSs). OCSs comprise fused input and output domains within a single regulatory protein, whereas TCSs consist of a sensor histidine kinase (HK) and response regulator (RR) that mediate diverse adaptive responses (Wadach et al., 2025). Quorum sensing enables population-dependent coordination of collective behaviours through extracellular autoinducers (Mukherjee & Bassler, 2019). In contrast to prototypical TCSs and quorum sensing, CSSs are multi-protein assemblies comprising 5-11 Che (<u>che</u>mosensory) proteins that regulate chemotaxis and other alternative cellular processes, including biofilm formation, type IV pili-mediated motility, development, virulence, and host interactions (Hickman et al., 2005; Munar-palmer et al., 2024). Their core components include the histidine kinase CheA, methyl-accepting chemotaxis proteins (MCPs), the response regulator CheY, and the scaffolding protein CheW, while CheR, CheB, CheD, CheV, CheC, CheZ, and CheX function as adaptation or auxiliary proteins. MCPs are membrane-bound or cytoplasmic receptors that detect extracellular or intracellular signals and relay them through CheW to CheA, which phosphorylates CheY to regulate downstream targets. In canonical chemotaxis pathways, phosphorylated CheY interacts with the flagellar switch proteins FliM and FliN to alter flagellar rotation and swimming behaviour (Ringgaard et al., 2018).

Based on comparative analyses of CSS genes’ organization, domain architecture, and inferred biological roles, bacterial CSS have been classified into flagellar (F), type IV pili-associated (Tfp), and alternative cellular function (ACF) classes. Flagellar classes have been categorised into 17 diversified classes (F1-F17), and other two groups contain a single class (Wuichet & Zhulin, 2010). The chemosensory systems have been studied in diverse bacterial species, such as *Escherichia coli* (Baker et al., 2006), *Pseudomonas aeruginosa* PAO1 (Ortega et al., 2017), *Vibrio cholerae* (Ortega et al., 2020)*, Myxococcus xanthus* (Sharma et al., 2018), etc. *Vibrio cholerae* possesses unusually complex chemosensory repertoires encoding three distinct chemosensory clusters, F6 and F9 on chromosome I and F7 on chromosome II. Cluster II/F6 forms the Short Membrane Arrays (SMA)-based large clusters at the poles of bacteria (flagellated poles) in proximity to flagellar genes (Ortega et al., 2020), and is well-known to regulate flagellar motility (Gosink et al., 2002; Hiremath et al., 2015; Hyakutake et al., 2005). Cluster I/F9 is purely cytoplasmic and has a characteristic double-layered appearance, whereas Cluster III/F7 is a long membrane array (LMA) (Ortega et al., 2020), and their functions remain incompletely understood.

Flagellar motility in *V. cholerae* is regulated by F6 CSS, which further helps in its colonization of the host intestine. *V. cholerae* shows chemotaxis response towards twenty L-amino acids, which is stronger than its response to carbohydrates (Boin et al., 2004). *V. cholerae* also shows chemotaxis response to chitin oligosaccharides, such as GlcNAc, a compound abundantly found in marine environments, indicating *Vibrio* species sense the surroundings through the F6 CSS cluster (Li & Roseman, 2004). *V. cholerae* motility has been proven crucial for epithelial colonization by penetrating the mucus layer (Almagro-Moreno et al., 2015), and it facilitates its dissemination throughout the proximal gut but not the distal part of the lumen (Almagro-Moreno et al., 2015). Unlike F6, F9 cluster forms cytoplasmic arrays in *V. cholerae* with the help of two signaling domains of its associated chemoreceptor DosM. F9 CSS genes are expressed mainly during stress conditions such as carbon starvation, low-oxygen conditions, and in late-stationary phase (Briegel et al., 2016; Hiremath et al., 2015). Similarly, cluster III/ F7 CSS components are expressed during the major stress-related sigma factor RpoS inside hosts (during infection), in stationary phase, and during carbon starvation (Briegel et al., 2016). Amongst other CSS components, MCPs play an integral role in communicating the signals to respective CSS. Within *V*. *cholerae* EI Tor N16961, only four MCPs, i.e., Mlp7, Mlp8, Mlp24, and Mlp30, have been reported to play a role in pathogenicity, whereas *mlp2*, *mlp29,* and *mlp42* genes are expressed during human infection (Matilla & Krell, 2018), signifying that motility and chemotaxis directly or indirectly contribute to host colonization and environmental adaptation.

Despite extensive experimental characterization of *V. cholerae* chemosensory systems, the diversity, distribution, and evolutionary relationships of these systems across the order Vibrionales remain poorly resolved. To address this gap, we performed a large-scale comparative genomic and phylogenetic investigation of signal-transduction systems across Vibrionales, both on the available complete genomes and metagenome-assembled genomes (MAGs). Using genome-wide homology detection, domain analysis, and structural bioinformatics, this study explores the diversity, distribution, and evolution of chemosensory systems and associated chemoreceptors across members of this ecologically and medically important bacterial order.

## Methodology

### Bacterial genomes under study

Genome data of 116 reference and representative Vibrionales organisms out of 7,250 available genome assemblies were downloaded from the NCBI RefSeq database on 10th February 2024, representing high-quality genomes across eight genera (Figure 1A). The host-association and habitat information was collected from literature studies and available metadata (Figure 1B, Table S1). Correlation analyses were performed using the cor.text function in R. We performed correlation analysis for genome sizes of both chromosomes and their respective CDS counts, genome size of large and small chromosomes, MCP protein count and genome size, total ribosomal RNA count and genome size, total tRNA count and genome size, orphan histidine kinase and chromosome genome size, orphan response regulator and chromosome genome size, and TCS count and chromosome genome size. The statistical significance between replicon genome size and GC content across different replicons, chromosome large, chromosome small, and plasmid were assessed using the ggsignif package of R.

**Figure 1.**
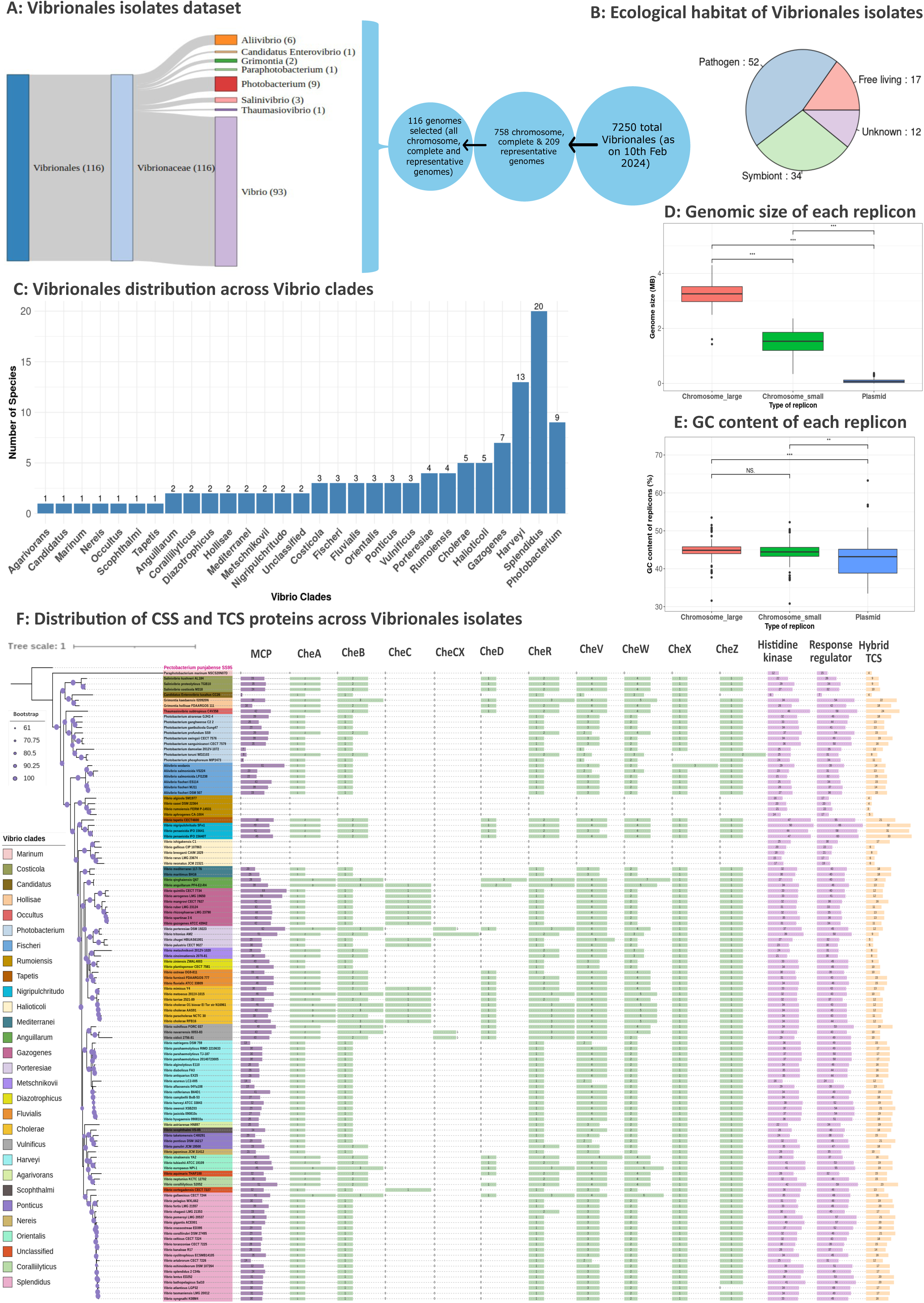
Genomic features and phylogenetic distribution of Vibrionales isolates used in this study. **(A)** Overview of selected 116 Vibrionales isolates included in this comparative genomic analysis. **(B)** Pie chart illustrates the diverse ecological habitats of Vibrionales members. **(C)** Distribution of isolates across twenty-eight different clades. **(D, E)** Box plots showing the distribution of genome size and GC content, respectively, across distinct replicon types. Statistical significance was assessed using the Pearson correlation coefficient test followed by pairwise comparisons; The significant P value is indicated in stars, *** indicates P < 0.001 (very highly statistically significant), ** indicates P < 0.01 (highly statistically significant), and NS indicates non-significant. **(F)** Genome-scale distribution of CSS proteins, orphan histidine kinases, orphan response regulators, and hybrid two-component system (TCS) proteins mapped onto the core-genome phylogeny using iTOL.

### Construction of core-orthologous gene-based phylogeny

Pan-genome analysis was performed for all Vibrionales along with an outgroup, *Pectobacterium panjabsese* SS95, using Proteinortho v6.0.14 (Klemm et al., 2023). The core, accessory, and unique genes across Vibrionales were identified using a custom shell script. Thirty genes out of 384 single-copy core genes were extracted, followed by their alignment using MUSCLE v5.1 (Edgar, 2022). Their concatenated alignment file of 12649 nucleotide sites was subjected to IQ-TREE 2.1.2 (Minh et al., 2020) to create a maximum likelihood phylogeny utilizing the best model (LG+F+R10 identified using IQ-TREE 2.1.2), with 1000 bootstraps, and the final tree was visualized in iTOL v6 (Letunic & Bork, 2024).

### Domain-based homology detection and identification of chemosensory clusters

Domains known to be associated with each CSS protein type were extracted from the Microbial Signal Transduction Database (MiST4.0) (Gumerov et al., 2024) and associated studies (Wuichet & Zhulin, 2010). The proteome of all Vibrionaceae members was scanned against Pfam-A v35.0 database with an E-value threshold of 1e^-5^ using hmmscan module of HMMER version 3.3.2 (Finn et al., 2011), using default parameters in domtblout format. The output was parsed and sorted using a shell script to avoid redundant overlapping domains. A total of 154 CheA proteins were categorized into different CheA-types by scanning these sequences against CheA-hmm profiles (Alexander & Zhulin, 2007) utilizing the hmmscan module of HMMER version 3.3.2 using above-mentioned parameters. Based on the location of all identified CSS proteins, the chemosensory clusters were identified from the GBFF files. The figure depicting the location of both F6 CSS clusters in *Photobacterium toruni* WD2103 was constructed using DNAPlotter (Carver et al., 2009).

### Construction of CheA gene phylogeny

154 CheA proteins from order Vibrionales and one CheA from outgroup *Pectobacterium punjabense* SS95 were aligned using MUSCLE v5.1, which were subjected to IQ-TREE 2.1.2 to generate a maximum likelihood phylogeny utilizing the best model, LG+F+R8 identified using IQ-TREE 2.1.2, with 1000 bootstraps and visualized in iTOL v6. The protein organization of each CSS was mapped onto the CheA phylogeny using iTOL v6.

### MCP protein identification and classification

Proteins having modular architecture with the marker domain of MCP protein [MCPsignal; PF00015.25] were extracted using a custom shell script to identify MCP proteins. The heptad classes of MCP proteins were identified by scanning all the MCP protein sequences against MCP-hmm profiles (Alexander & Zhulin, 2007) using the hmmscan module of HMMER version 3.3.2 using above parameters, and their distribution was mapped on core gene phylogeny. A box plot illustrating the distribution of MCP heptad types per replicon, encompassing both chromosomes and plasmids, was generated using the ggplot2 package in R. A pie chart was constructed to depict the distribution of diverse ligand-binding domains (LBDs) of the MCP protein using the plotrix, dplyr, and RColorBrewer packages in R. Forty six sequences of *Vibrio cholerae* O1 biovar EI Tor N16961 MCPs were aligned using MUSCLE v5.1, which was subjected to IQ-TREE 2.1.2 to generate a maximum likelihood phylogeny utilizing the best model from IQ-TREE 2.1.2, LG+F+R5, with 1000 bootstraps and domain architecture were mapped using iTOL v6.

### Sequence network analysis of CheA proteins

154 CheA proteins were clustered using MMseqs2 18.8cc5c (Steinegger & Söding, 2017) with a threshold of minimum sequence identity of 0.4 and alignment coverage of 0.8, which was visualized in Cytoscape 3.10.3 (Shannon et al., 2003).

### CheA structures download, prediction, alignment and comparison

25 out of 154 CheA structures, available on AlphaFold database, were downloaded using AlphFoldStructureExtractor tool (Saraf et al., 2025). CheA-F9 structure was not available on the AlphaFold database; therefore, CheA-F9 protein structure was predicted using the AlphaFold server (https://alphafoldserver.com/), and output CIF file format was converted to PDB format using PyMOL v2.6.2 (The PyMOL Molecular Graphics System, Version 2.6.2 Schrödinger, LLC.) for further visualization and structural alignment. For structures validation, we have used the pLDDT and pTM score for already available and predicted structures, respectively. Representative structure from each CheA class was aligned to each other using TM-align (Zhang & Skolnick, 2005) to calculate TM-score and RMSD score.

### CheA homologs phylogeny construction

All 154 CheA sequences were subjected to diamond blastp v2.1.0.154 (McGinnis & Madden, 2004) against the non-redundant (NR) database with an E-value threshold of 1e^-5^. Top fifty non-Vibrionaceae hits for each CheA protein were subjected to multiple sequence alignment using MUSCLE v5.1 (Edgar, 2022). The alignment was subjected to FastTree v2.1.11 (Price et al., 2009) to generate a maximum likelihood phylogeny that was visualized in iTOL v6 (Letunic & Bork, 2024). Top 50 non-Vibrionaceae hits were separately extracted for F9 CheA protein sequence, which were aligned using MUSCLE v5.1, and aligned sequences were subjected to IQ-TREE 2.1.2 to get a maximum likelihood phylogeny utilizing the best model, LG+F+R7 identified using IQ-TREE 2.1.2, with 1000 bootstraps, and finally visualized in iTOL v6. The protein architecture of each class Alphaproteobacteria F9 CheA homolog was identified from their genomes and depicted on the phylogeny along with mapping of the order and class-level taxonomy using iTOL v6.

### Identification of CheA from ∼10,000 order Vibrionales RefSeq genomes and MAGs

All available (9,667 genomes and 159 MAGs) genomes were downloaded from NCBI in August 2025. CheA proteins along with other CSS proteins were identified and classified into respective CSS types as mentioned above. All CheA proteins were clustered using CD-HIT 4.8.1 (Fu et al., 2012) with a threshold of 0.9 sequence identity. Clustered CheA sequences, along with the outgroup, were aligned using MUSCLE v5.1, which was subjected to FastTree v2.1.11 to generate a maximum likelihood tree, and visualized using iTOL v6, where identified CheA classes and number of CheA proteins clustered in each representative CheA protein were mapped on phylogeny.

### BLASTn to identify the duplicated region of the genome in *V. qinghaiensis* Q67

BLASTn 2.2.30 (McGinnis & Madden, 2004) analysis was performed to identify duplicated genomic regions in *V. qinghaiensis* Q67. A BLAST database was created using the genomic .fna file, which was also used as the query input.

## Results

### Comprehensive dataset captures phylogenetic, ecological, and genomic diversity across order Vibrionales

To understand the comprehensive diversity of signal transduction and CSS proteins, 116 order Vibrionales genomes from eight out of fifteen diverse genera (Figure 1A) and different ecological habitats (Figure 1B) were selected for this comparative study. These representative/reference and high-quality genomes have ∼80% (93/116), ∼7% (9/116), and ∼5% (5/116) representation from the *Vibrio*, *Photobacterium,* and *Aliivibrio* genera, respectively. Besides this, a few genera such as *Candidatus Enterovibrio*, *Grimontia*, *Paraphotobacterium*, *Salinivibrio*, and *Thaumasiovibrio* have only a few representatives (1-3) in our dataset. Seven known genera, i.e., *Allomonas*, *Allocatenococcus*, *Enterovibrio*, *Photococcus*, *Corallibacterium*, *Veronia*, and *Parasalinivibrio,* have only draft genomes, and owing to this, we have not included them in our dataset. Members of the order Vibrionales exhibit remarkable ecological versatility, encompassing a broad spectrum of lifestyle strategies ranging from mutualistic symbiosis to pathogenicity and free-living existence. In our dataset, 52, 34, and 17 species are pathogenic, symbiotic, and free-living, respectively (Figure 1B), and out of all species, Splendidus and Harveyi clades contain a larger number of species (Figure 1C). Several species establish symbiotic associations with marine hosts, exemplified by *Aliivibrio fischeri*, which forms a well-characterized mutualism with squid (Miyashiro & Ruby, 2012), contributing to bioluminescence-mediated host interactions. In contrast, some of the members of the Vibrionales order are known to demonstrate opportunistic and obligatory pathogenic potential to a wide range of animals, including marine animals such as shrimp, shellfish, and coral reefs, as well as to humans, where some of the members, such as *V. parahaemolyticus* and *V. cholerae,* cause vibriosis and cholera, respectively (Baker-Austin et al., 2018). Additionally, certain members, such as *V. vulnificus* MO6-24/O, have been shown to infect plant hosts including *Arabidopsis thaliana* under artificial conditions, highlighting cross-kingdom pathogenic potential (Park et al., 2021). Alongside these host-associated lifestyles, a subset of Vibrionales persists as free-living organisms in aquatic environments, reflecting their ability to adapt to diverse ecological niches. Apart from diverse ecological habitats, Vibrionales members possess a characteristic bipartite genome, consisting of two chromosomes and, in some species, additional plasmids. Chromosome I is typically larger in genomic size (average 3.25 Mb) as compared to chromosome II (average of 1.53 Mb) (Figure 1D). Irrespective of different genomic sizes, the GC content for both chromosomes is the same (Figure 1E). Considerable variation was also observed in rRNA and tRNA gene copy numbers across different lineages, reflecting the genomic diversity within the order (Figure S1). Core-gene phylogeny generated using 30 housekeeping genes was able to demarcate all Vibrionales members across their 28 *Vibrio* clades as per the latest clade classification (Jiang et al., 2022).

### Vibrionales organisms exhibit extensive diversification in signal transduction components and chemosensory proteins

Order Vibrionales members encode a highly heterogeneous and expanded repertoire of signal transduction systems, including hybrid TCSs, CSSs, and orphan signaling proteins (Figure 1F). Orphan histidine kinases and response regulators, as defined by signaling proteins lacking association with known CSSs, are ubiquitously distributed across Vibrionales genomes, with average values of 31 and 41 (median of 32 and 43), respectively, showcasing the potential modularity and rewiring capacity of these signaling architectures. Notably, hybrid TCS proteins (average and median of 15), harboring both kinase and receiver domains within a single polypeptide, are less abundant than HK and RR, suggesting an increased propensity for multistep phosphorelay mechanisms and enhanced signal integration. TCS protein count varies between different species, ranging from three in *V. rumoiensis* FERM P-14531 to 33 in *V. penaeicida* IFO 15640T, suggesting lineage-specific expansion of the repertoire of TCS proteins and differential ecological tuning of signaling repertoires. On the other hand, correlation analysis revealed no statistically significant association between chromosome size and the abundance of orphan HKs, orphan RRs, or hybrid TCS proteins (p ∼ 0.7; Figure S2), suggesting that the diversification and expansion of the TCS proteins are not simply a function of genome size scaling. Instead, these patterns reflect niche-specific selective pressures and adaptive signaling requirements. Furthermore, clade-level data indicate that the Marinum, Rumoiensis, and Halioticoli clades encode comparatively fewer hybrid TCS proteins relative to other *Vibrio* clades. This trend points toward a potential functional linkage or co-evolutionary constraint between CSS complexity and hybrid TCS abundance, implying that expansion of chemosensory associated signaling may be coupled with increased reliance on hybrid phosphorelay architecture.

### Chemosensory system distribution correlates with motility loss in specific Vibrio clades

Comparative genomic analysis across Vibrionales members reveals that the core CSS components (CheA, CheW, CheY, CheB, and CheR) are broadly conserved across most clades, while MCP abundance varies sporadically among different species/clades. Such conservation of core proteins putatively suggests the evolutionary maintenance of chemosensory signaling capabilities, which highlights their importance in environmental sensing and motility. In contrast, accessory proteins (e.g., CheC, CheCX, and CheD) exhibit a patchy phylogenetic distribution and are absent in several lineages, suggesting lineage-specific functional specialization or redundancy (Figure 1F). Replicon-level distribution indicates a strong genomic bias in the localization of CSS genes, as most chemosensory components are encoded on chromosome I, whereas chromosome II harbors fewer CSS-related genes, and plasmids encode only a minimal and sporadic subset (Table S3). This pattern suggests that chromosome I serves as the primary genomic scaffold for core signal transduction pathways, while the presence of these CSS components on secondary replicons may contribute auxiliary or niche-specific adaptations for specific organisms within specific clades.

The counts of an important CSS marker gene, *CheA,* which indicates the number of CSS clusters encoded by species, range from 1 to 3 across all *Vibrio* clades (Figure S3A). As compared to other CSS proteins, MCP proteins are significantly higher in all Vibrionales members, which range from 2 to 64 (median of 33), lowest *in Candidatus Enterovibrio luxaltus* CC26 (Candidatus clade) and highest in *V. quintilis* CECT 7734 (Gazogenes clade). Overall, species within the Gazogenes clade, along with some species from Cholerae, Coralliilyticus, Diazotrophicus, Fluvialis and many other clades possess a greater number of MCP proteins (greater than average MCP count of 33), potentially linked to diverse stimuli received by these members from their habitat (Figure S3B, Table S6). Our study identified that *V. cholerae* O1 biovar El Tor strain N16961 encodes a total of 46 MCP proteins and 22 other chemosensory proteins, which is consistent with the previous study (Boin et al., 2004) (Table S4). The CheA count remains relatively conserved across different *Vibrio* clades, indicating limited variation across the clade, suggesting its evolutionarily conserved need and nature (Figure S3A). The distribution of MCP counts across *Vibrio* clades reveals substantial variability, suggesting that this diversity is shaped by the ecological niche and habitat of these members (Figure S3B).

All core chemosensory system (CSS) components are broadly conserved across the examined clades, with notable exceptions in the species from non-motile Marinum, Rumoiensis, and Halioticoli clades. This pattern supports a model of coordinated evolutionary reduction, wherein the loss of motility is accompanied by the concomitant degeneration or complete loss of associated CSS modules, including those directly implicated in flagellar regulation. Such co-elimination of CSS and flagellar genes likely reflects selective pressures favoring genome streamlining and the dispensability of chemotaxis in specific ecological niches (Geng et al., 2020; Hidalgo et al., 2009; Huang et al., 2016).

### The F8 chemosensory system represents an experimentally unexplored system in Vibrionales

CheA homologs per genome strongly correlate with the number of CSS clusters encoded, consistent with the modular organization of CSS pathways. Therefore, to systematically characterize the diversity of Vibrionales CSSs, a comprehensive phylogenetic analysis was performed using 154 identified CheA sequences. The resulting tree resolved into four well-supported and deeply branching clades, corresponding to four discrete CSS types, which include the previously described F6, F7, and F9 systems, along with an experimentally uncharacterized CSS system, F8 (Figure 2A). Integration of phylogenetic clustering with genomic context analysis revealed that each CheA-defined lineage is associated with a distinct and conserved gene cluster architecture, consisting of a unique combination of core and accessory che proteins (Figure 2B). The F8 CSS cluster constitutes core proteins including CheA, CheW, CheY, CheB, CheR, as well as accessory protein CheD. Some MCP proteins were also reported within or near this F8 CSS cluster (Figure 2A). This indicates the putative coupling of these MCP proteins to F8 CSS cluster. In addition to canonical chemosensory components, all four CSS clusters also encode non-chemosensory-associated proteins (annotated as AAA in Figure 2B), indicating potential functional coupling with other cellular processes. Of particular interest are highly conserved ParA family ATPases (ParC; part of ParC/ParP protein complex) within F6 clusters, which are known to mediate polar localization of chemosensory complexes/arrays by interacting with the HubP protein as an anchoring factor (Altinoglu et al., 2022; Ringgaard et al., 2011).

**Figure 2:**
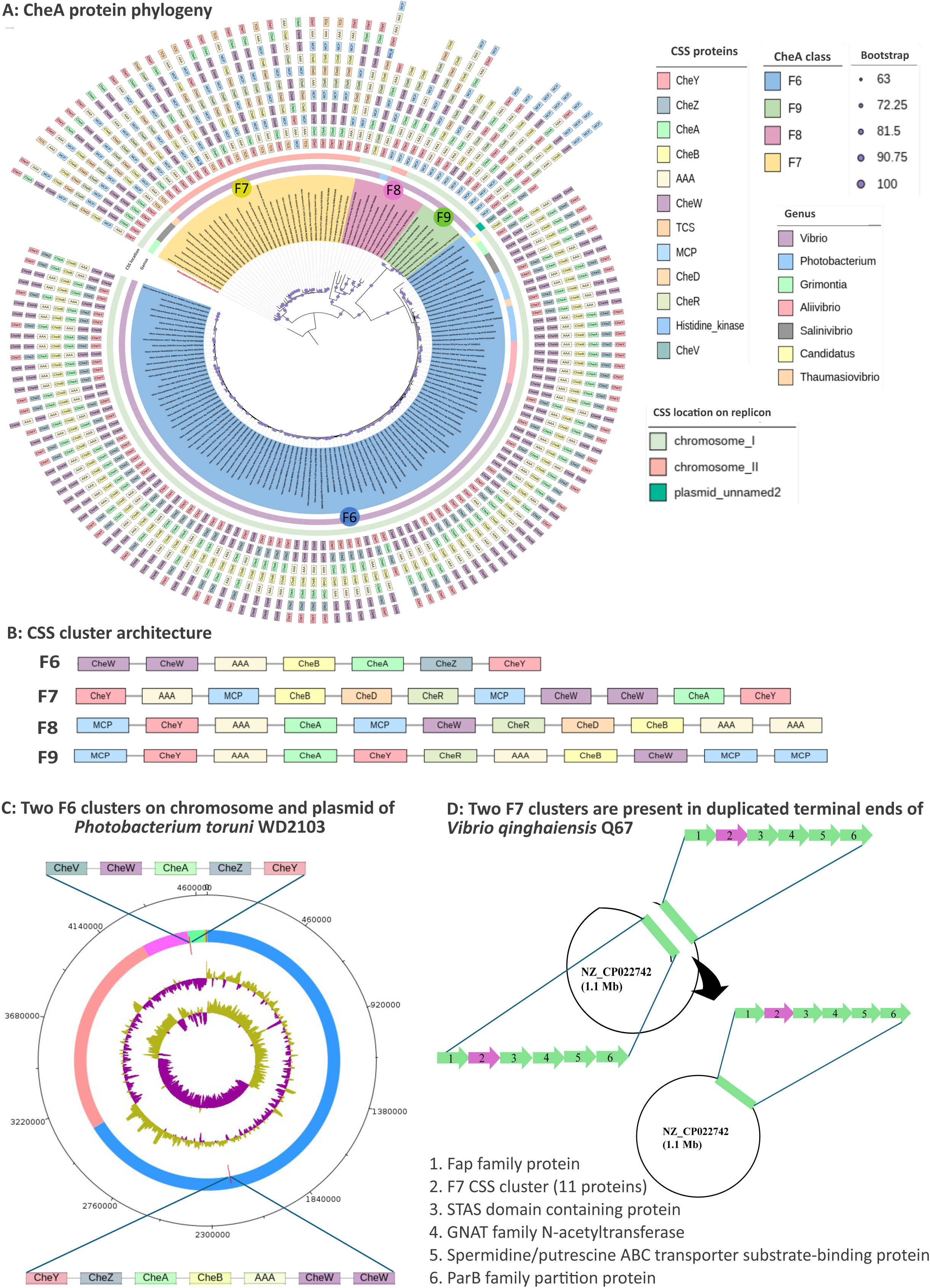
Distribution and genomic organization of CSS clusters in Vibrionales. **(A)** CheA-based maximum-likelihood phylogeny delineating into four distinct CSS clades (F6, F7, F8, and F9). All CheA proteins were aligned using MUSCLE v5.1 (block length of 1,429), for which phylogenetic inference was performed under the LG+F+R8 substitution model. This phylogeny was annotated with the CSS architecture types using iTOL. **(B)** Domain architectures of representative CSS clusters for each lineage. **(C)** Circos plot illustrating the replicon-level distribution of two F6 clusters in *Photobacterium toruni* WD2103. Replicons are color-coded as follows: chromosome I (blue), chromosome II (orange), plasmid 1 (pink), and plasmid 2 (light green). **(D)** Genomic context of two F7 clusters located within duplicated terminal regions of *V. qinghaiensis* Q67, indicating a potential segmental duplication while comparing the terminal ends of a chromosome.

Out of four identified CSS types (F6, F7, F8, and F9) (Figure 2A; Table S5), the F6 system is ubiquitous across Vibrionales and primarily associated with flagellar motility, suggesting its essential role in environmental sensing and motility. However, it is absent in *V. qinghaiensis* Q67 and a few additional non-motile clades, Marinum, Rumoiensis, and Halioticoli, advocating lineage-specific loss or functional replacement under particular ecological conditions. It must be noted that *V. qinghaiensis* Q67 has lost F6 but retained other CSS types; however, species from Marinum, Rumoiensis, and Halioticoli clades do not encode any CSS. The F7 system exhibits a more restricted but partially-clade-conserved distribution, being sporadically present in members of the Costicola (3/3 organisms), Hollisae (2/2 organisms), Nigripulchritudo (3/3 organisms), Anguillarum (2/2 organisms), Fluvialis (3/3 organisms), Cholerae (7/7organisms), Orientalis (3/3 organisms), Coralliilyticus (2/2 organisms), Occultus (1 organism), Splendidus (1/20 organisms), Tapetis (1 organism), and Vulnificus (1/3 organisms) clades. Earlier, the F9 has been reported in *V. cholerae* strains (Briegel et al., 2016); however, this comparative study identified F9 to be extremely rare, exhibiting its presence only in two *Vibrio* clades, i.e., Cholerae (5/7 organisms) and Porteresiae (2/4 organisms), indicating a highly specialized or condition-dependent role, which needs to be further explored to understand what ecological adaptations these organisms might need for having this CSS type.

Like F7, the F8 system also displays a patchy phylogenetic distribution, being detected in selected species belonging to Photobacterium (1/9 organisms), Anguillarum (2/2 organisms), Porteresiae (1/4 organisms), Metschnikovii (2/2 organisms), Vulnificus (2/3 organisms), Orientalis (1/3 organisms), and Splendidus (1/20 organisms) clades. The sporadic presence of F8 along with previously known F7 within specific ecological groups raises the possibility that this system mediates specialized sensory responses, potentially linked to host association, virulence, or niche adaptation, which needs to be explored further. An exception to this pattern is observed in *V. anguillarum* PF4-E2-R4, *V. qinghaiensis* Q67 (Anguillarum), *V. europaeus* NPI-1 (Orientalis), and *V. gallaecicus* CECT 7244 (Splendidus clade), which uniquely encode both F7 and F8 systems, unlike other members of the same clade. This atypical configuration may reflect recent acquisition events or selective retention of multiple CSS modules to enhance ecological versatility.

### Replicon-specific organization of CSS clusters links genome architecture to functional hierarchy

We further examined the genomic organization and distribution of CSS clusters across replicons within Vibrionales (Figure S4). CSS loci were distributed across both primary and secondary chromosomes, as well as plasmids, highlighting their genomic plasticity. In all *V. cholerae* strains, the F9 (cluster I) and F6 (cluster II) loci are localized on chromosome I; however, the F7 (cluster III) locus is localized on chromosome II, which is consistent with previous studies (Hiremath et al., 2015). Across Vibrionales, the F6 system is predominately localized on chromosome I, supporting its role as a core, conserved motility-associated system. A notable exception is observed in *Photobacterium toruni* WD2103, which harbours two F6 clusters, one located on chromosome I and the other on a plasmid. The plasmid-encoded F6 cluster lacks the CheB methylesterase, suggesting potential functional divergence or incomplete adaptation machinery (Figure 2C). The F7 system is preferentially localized on chromosome II in *Vibrio* species, whereas in non-*Vibrio* clades such as Costicola, Hollisae, and Occultus, it is found on chromosome I, suggesting lineage-specific genomic rearrangements and potential differences in regulatory integration. The F8 system exhibits a dual chromosomal distribution, with notable presence on chromosome II in the Vulnificus clade, supporting its classification as a potentially adaptive or accessory system. In contrast, F9 clusters are consistently located on chromosome I, indicating a more stable genomic integration.

We also report that *V. qinghaiensis* Q67 has two F7 CSS clusters, both located-on chromosome II. However, we found this to be an interesting case of a genome assembly artifact, resulting in two F7 systems. Detailed sequence analysis of the circular genome of *V. qinghaiensis* Q67 revealed that these clusters reside within identical (∼99% sequence identity) terminal regions spanning ∼20.3 kb at both ends of the replicon. This high degree of similarity on both terminals indicates an assembly artifact involving duplicated terminal sequences (Figure 2D). After accounting for this duplication, only a single F7 cluster is retained in the corrected genome model. This observation is consistent with similar cases reported in other bacteria, *Myxococcus xanthus* DZ2, and underscores the importance of careful validation of repetitive genomic regions when interpreting gene copy number and system multiplicity (Mahanta & Sharma, 2025).

Taken together, these results suggest replicon-specific partitioning of CSS clusters, with core system (F6) showing genomic conservation, whereas accessory systems (F7, F8, and F9) exhibit distributional flexibility. Such patterns are putatively consistent with a functional hierarchy in which conserved system support motility while other CSS classes may contribute to ecological adaptation.

### Large-scale study of >10,000 Vibrionales genomes and MAGs further support the presence of only four chemosensory systems

To further comprehensively assess the diversity and distribution of CSS across the order Vibrionales, we also performed a high-throughput large-scale comparative analysis using the histidine kinase CheA as a conserved phylogenetic marker. A total of 14,416 CheA homologs were identified from 9,667 RefSeq genomes and 159 MAGs, representing one of the most extensive datasets compiled to date for chemosensory system characterization. To reduce redundancy and capture representative sequence diversity, CheA homologs were clustered using CD-HIT, resulting in 329 non-redundant clusters. Phylogenetic reconstruction of these representative sequences further revealed the presence of only four well-supported clades corresponding to F6, F7, F8, and F9 CSS types (Figure 3), as identified with the smaller 116-organism dataset (Figure 2A), indicating that they constitute the whole chemosensory repertoire of Vibrionales.

**Figure 3:**
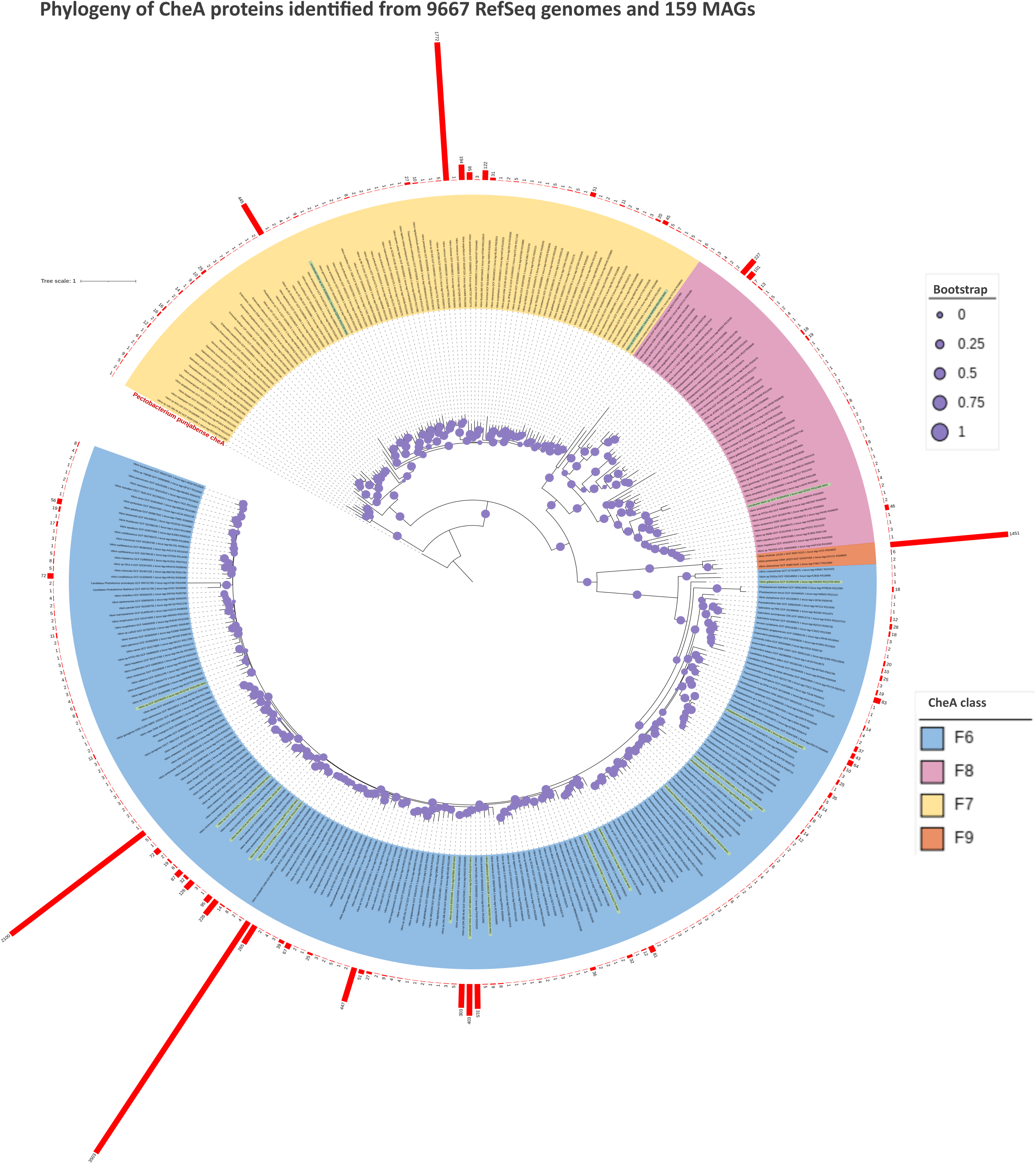
Maximum-likelihood phylogeny of CheA proteins from ∼10,000 order Vibrionales genomes and metagenome-assembled genomes (MAGs). All clustered CheA proteins were aligned using MUSCLE v5.1, and their final alignment block length of 2,223 was used for this amino acid model-based ML phylogeny. The four CSS lineages (F6, F7, F8, and F9) are highlighted in distinct colours. MAG-derived sequences are indicated in green. The bar plot adjacent to the phylogeny denotes the number of CheA sequences assigned to each lineage, illustrating the relative representation of each CSS type across the dataset.

In contrast to the widespread occurrence of CSS in RefSeq genomes, their representation in MAGs was markedly reduced. Among the 159 MAGs analysed, only eighteen encoded identifiable CSS clusters, suggesting incomplete genome reconstruction. Within these MAGs, F6 was the most prevalent system, detected in fifteen genomes, consistent with its established role in flagellar motility and its near-universal presence across Vibrionales. F7 was identified in only two MAGs, while the F8 system was detected in a single MAG, highlighting its low abundance and potentially specialized functional role. Notably, the F9 system was absent from all MAGs, suggesting that it may be associated with specific ecological niches, host-associated lifestyles, or genomic contexts that are underrepresented in metagenomic assemblies. Alternatively, the absence of F9 or other CSS types in MAGs may also reflect assembly limitations, as chemosensory arrays are often organized as large clusters that are difficult to recover from fragmented metagenomic data.

### SSN and comparative structural analyses reveal modular diversification of CheA proteins within a conserved chemosensory signaling scaffold

Applying SSN analysis to the Vibrionales CheA dataset at stringent thresholds (≥40% sequence identity and ≥0.8 alignment coverage) resolved the 154 CheA homologs into four well-defined network communities (Figure 4A), corresponding to the phylogenetically inferred F6, F7, F8, and F9 classes (Figure 2A). Notably, a single CheA homolog from *V. porteresiae* DSM 19223 (F8-type) formed an isolated node, failing to cluster with other F8 members, indicative of extreme sequence divergence. This observation is concordant with phylogenetic reconstruction, wherein this protein branches independently from all other CheA lineages, suggesting sequence divergence. The lack of detectable sequence similarity (<40%) with other CheA homologs potentially represents a transitional or highly specialized variant within the F8 lineage. In contrast, proteins within each SSN-defined cluster exhibit ≥40% identity, consistent with established thresholds for structural conservation and functional equivalence.

**Figure 4:**
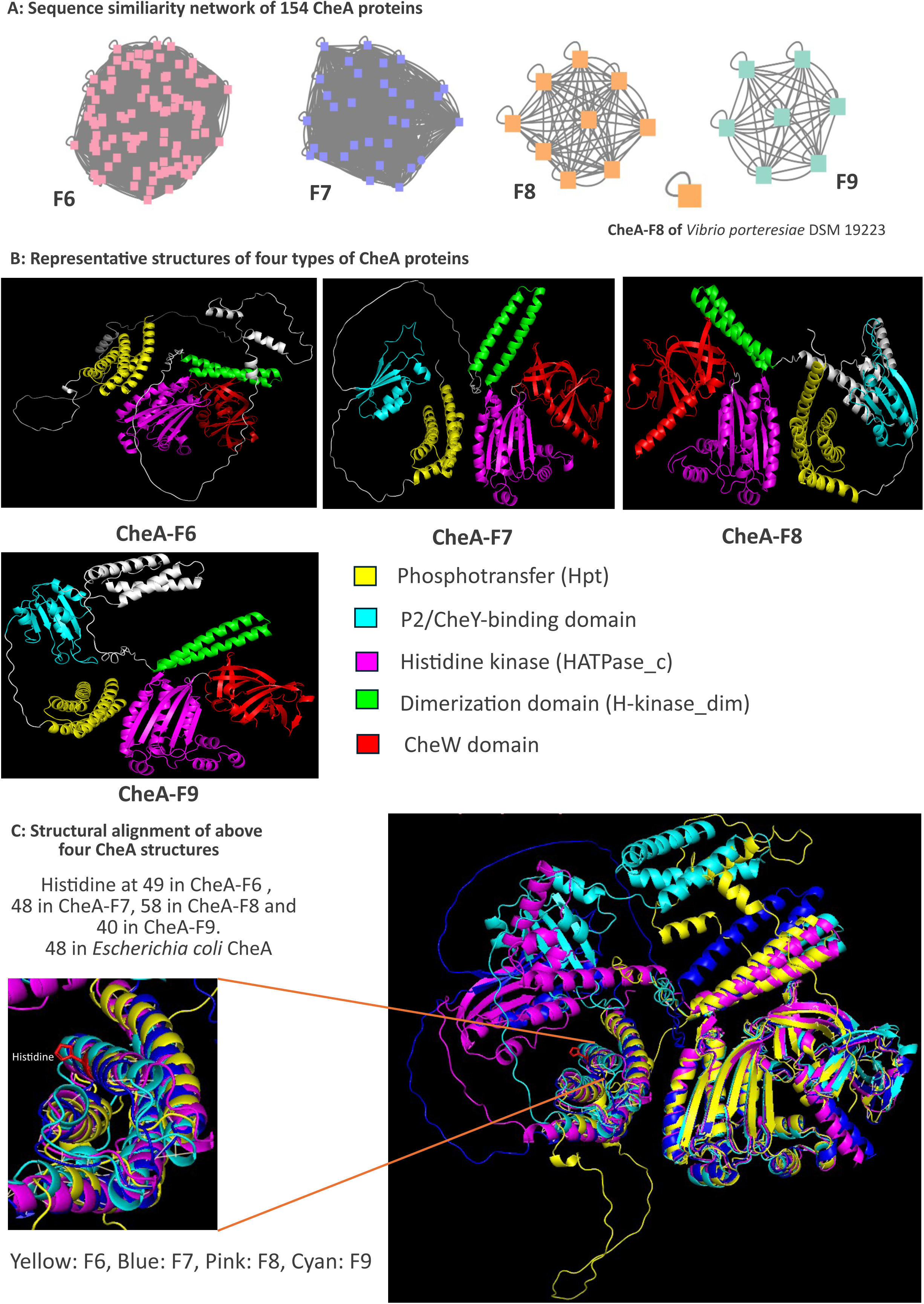
Structural characterization of CheA proteins across Vibrionales CSS lineages. **(A)** Sequence similarity network (SSN) analysis of CheA proteins, revealing four discrete clusters corresponding to CSS types F6, F7, F8, and F9; a notable exception was identified within the F8 cluster, indicating potential structural divergence within this lineage. **(B)** Four panels showing the domain architecture of representative CheA proteins from each CSS type, with the five conserved functional domains highlighted in distinct colours. CheA-F6: *Aliivibrio wodanis* (locus tag= AWOD_RS10085, AlphaFold ID:AF-A0A090IUF0-F1), CheA-F7: *Vibrio aquimaris* THAF100 (locus tag=FIV01_RS18245, AlphaFold ID: AF-A0A5P9CQY6-F1), CheA-F8: *Vibrio qinghaiensis* Q67 (locus tag=CCZ37_RS06945, AlphaFold ID:AF-A0A223MXP1-F1), and CheA-F9: *Vibrio cholerae* O1 biovar El Tor str N16961 (locus tag=FY484_RS07030). The structure of CheA-F9 was predicted using the AlphaFold server (https://alphafoldserver.com/). **(C)** Structural superimposition of representative CheA proteins from each lineage, demonstrating conservation of the catalytic histidine residue within the phosphotransfer (Hpt) N-terminal domain across all CheA types.

To further question the structural basis of this diversification, high-confidence models of representative CheA structures for each class were extracted from the AlphaFold Protein Structure Database (Figure 4B). Canonical CheA proteins are characterized by a conserved five-domain architecture, comprising the P1 (Hpt/phosphotransfer), P2 (CheY-binding), P3 (dimerization; H-kinase_dim), P4 (ATP-binding; HATPase_c), and P5 (CheW-like regulatory) domains, as established in model systems such as *Escherichia coli*, *Thermotoga maritima*, *Salmonella enterica*, *Bacillus subtilis*, and *Halobacterium salinarum*. Previous reports have suggested that only 46% of CheA protein sequences have this P2 domain; therefore, only P1, P3, P4, and P5 should be considered as key domains for CheA protein (Berry et al., 2024). Prior biochemical studies have also demonstrated that the P2 domain, while enhancing phosphotransfer efficiency, is not strictly essential for CheA-mediated signaling, and its functional contribution can be partially compensated by increased Hpt (P1) activity (Jahreis et al., 2004). Structural comparisons revealed that representative CheA-F7 retains this canonical five-domain organization and closely resembles the archetypal CheA architecture of *E. coli*, supporting its conserved role in chemosensory signaling (Figure 4B). However, in our predicted CheA-F6 structures, the P2 domain was not resolved as a distinct structural unit, which is likely attributable to intrinsic disorder or limitations in structure prediction rather than true domain loss, as supported by extended low-confidence regions between P1 and P3, as supported by previous studies (Berry et al., 2024).

Noticeably, structural models of representative CheA-F8 and CheA-F9 proteins revealed an additional insertion between the predicted P2 and P3 domains. Despite these variations, structural superposition analysis revealed that the catalytic histidine residue within the P1 domain, corresponding to His48 in *E. coli*, is strictly conserved across all CheA types (Figure 4C), showcasing strong evolutionary constraints on phosphotransfer functionality (Muok et al., 2020). Since RMSD weights the distances between all residue pairs equally, a small number of local structural deviations could result in a high RMSD, even when the global topologies of the compared structures are similar (Zhang & Skolnick, 2005). The Template Modelling score (TM-score) is independent of protein sequence length and considers the global topology of the protein structure. Despite pronounced sequence divergence and SSN-based segregation, all pairwise structural comparisons yielded TM-scores in the range of 0.6-0.7, indicative of a conserved global fold across all CheA types (Figure S5). This apparent discrepancy between sequence and structural divergence highlights that CheA proteins have undergone extensive sequence diversification while preserving a conserved three-dimensional scaffold for kinase activity and signal transduction. Collectively, these findings suggest that CheA evolution in Vibrionales is characterized by modular diversification, domain insertions, and lineage-specific adaptations, while maintaining a structurally conserved core necessary for chemosensory signaling.

### Horizontal acquisition of the F9 chemosensory system from Alphaproteobacteria shapes Vibrionales signaling diversity

To understand the putative evolutionary origins of Vibrionales CSSs, a high-throughput maximum-likelihood phylogeny (Figure 5A) based on the top fifty non-Vibrionaceae CheA homologs robustly resolved four deeply branching clades corresponding to four CSS types, with Vibrionales sequences forming monophyletic or near-monophyletic subclusters within each clade. Notably, F6 and F7 systems were broadly distributed across Gammaproteobacteria, with dense representation among Vibrionales, consistent with prior studies indicating that these systems constitute ancestral, vertically inherited chemosensory modules that have been retained since early diversification of this lineage. In contrast, the F8 lineage, although detected across phylogenetically distant classes including Desulfovibrionia, Desulfobulbia, and Desulforomonadia, exhibited its highest prevalence within Gammaproteobacteria, suggesting that these groups also likely represent the ancestral reservoir of F8 systems across Gammaproteobacteria, followed by lineage-specific retention within a few selected Vibrionales dependent on their ecological dependencies.

**Figure 5:**
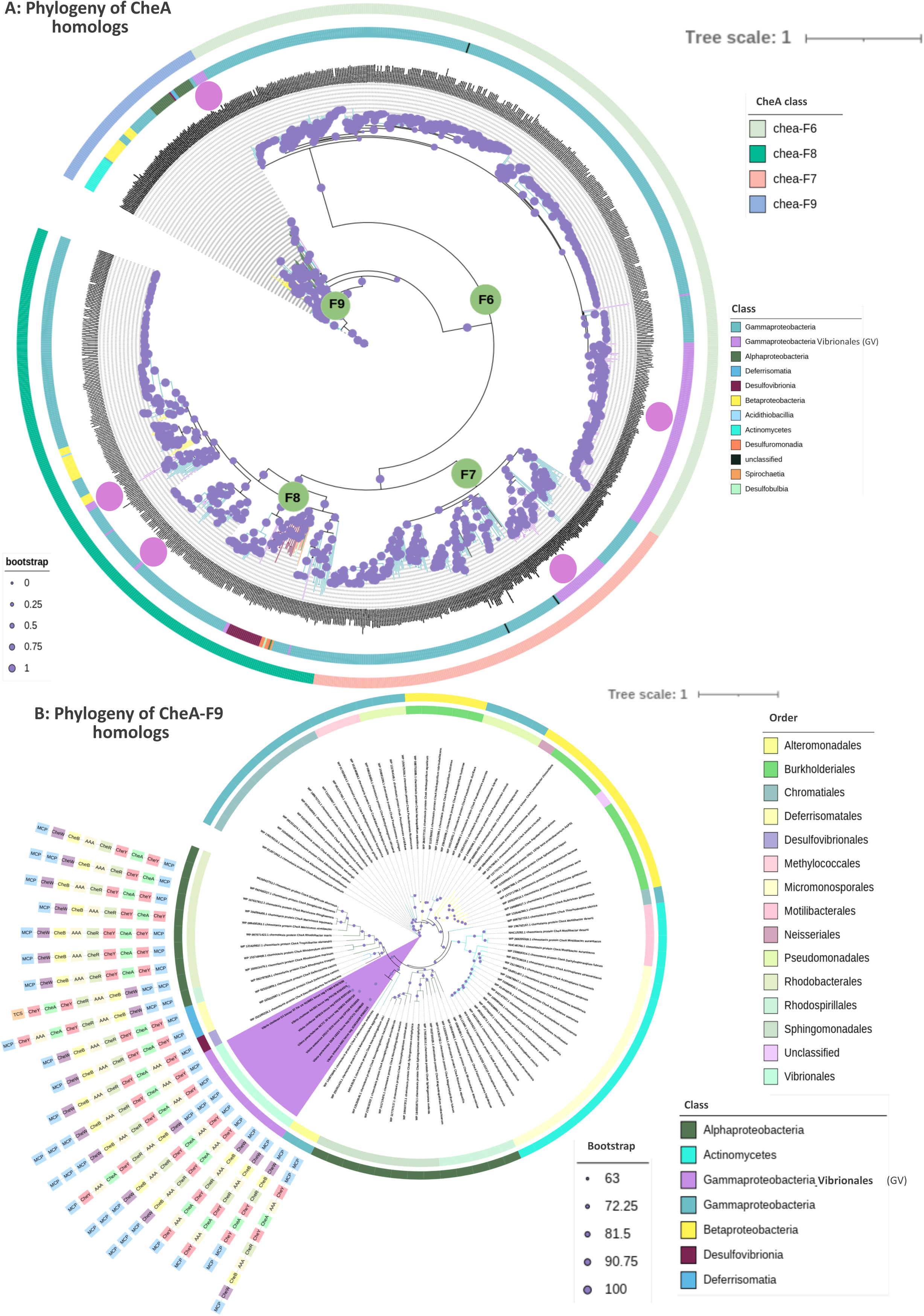
Phylogenetic analysis of CheA homologs with emphasis on the evolutionary origin of the F9 CSS lineage. **(A)** CheA homolog phylogeny demarcated the phylogeny into four distinct CSS clusters, namely F6, F7, F8, and F9. The outer ring indicates the CSS cluster type, while the inner ring circle represents class-level taxonomy. All CheA homologs along with CheA proteins were aligned together using MUSCLE v5.1, and their final alignment block length of 2,743 was used for this amino acid model-based ML phylogeny. This phylogeny was visualized in iTOL. **(B)** Focused phylogeny of F9 CheA homologs, annotated with CSS cluster domain architectures from representative classes including Alphaproteobacteria, Desulfovibrionia, Deferrisomatia, and Gammaproteobacteria (Vibrionales). The outer and inner rings indicate class and order, respectively. The purple-coloured clade represents the Gammaproteobacteria_Vibrionales in the phylogeny. The sequences were aligned using MUSCLE v5.1, and the alignment block length of 1,285 was used for this LG+F+R7 model-based ML phylogeny and visualized in iTOL.

Strikingly, F9-associated CheA homologs displayed a broad but discontinuous distribution spanning Gammaproteobacteria, Alphaproteobacteria, and Actinomycetota, with Vibrionales sequences phylogenetically embedded within or closely affiliated to Alphaproteobacterial clades (Figure 5B). This observation supports a putative horizontal acquisition of the F9 system by Vibrionales from an Alphaproteobacterial lineage. Such inter-class HGT events have been increasingly recognized as a major force shaping the evolution of bacterial sensory systems, particularly in ecologically versatile taxa such as *Vibrio* that frequently encounter fluctuating environmental and host-associated niches. Further supporting this hypothesis, comparative analysis of protein architecture of F9 CSS revealed a high degree of similarity between Vibrionales F9 protein architecture and their Alphaproteobacterial counterparts, including preservation of key histidine kinase domains, receiver modules, and associated regulatory components (Figure 5B). This architectural resemblance, coupled with phylogenetic clustering, suggests not only a shared evolutionary origin but also potential conservation of signaling mechanisms and functional outputs.

Collectively, these analyses suggest that different CSS classes may have followed distinct evolutionary trajectories within Vibrionales. Amongst the four CSS, F6 and F7 appear broadly conserved across the order Vibrionales and are putatively inherited from nearby Gammaproteobacteria; F8 showcases sporadic presence in a few Vibrionales but is quite dominant across Gammaproteobacteria; however, the F9 have likely been acquired via horizontal gene transfer, potentially contributing to the sensory diversification and ecological adaptation of order Vibrionales.

### The diversity in ligand-binding domains and heptad classification of MCP indicate its central role in the chemosensory system

Comprehensive identification of MCPs across 116 Vibrionales genomes revealed a large repertoire of MCPs (ranging from 2 to 64 per genome; median value of 33); however, only a few MCPs are genomically associated with CSS, whereas the vast majority are dispersed across the genome and not even present near the CSS (Figure 2B). Species within the Gazogenes and Cholerae clades encode a higher number of MCPs, likely reflecting their diverse ecological niches; for example, *V. cholerae* inhabits complex gut environments where it senses bile and intestinal mucin (O’Toole et al., 1999). Notably, cluster-associated MCPs were restricted to F7-, F8-, and F9-type systems, whereas F6 systems consistently lacked proximal MCP-encoding genes in all the organisms without any exception, suggesting that F6 modules may function as core signaling backbones that integrate inputs from different MCPs, thereby enhancing signal integration capacity.

To further resolve MCP diversity, these proteins were classified based on the length of their cytoplasmic coiled-coil regions, defined by heptad repeat architecture (Figure 6A, 6D) (7-residue periodicity), which determines receptor packing geometry and signaling state transitions (Alexander & Zhulin, 2007). The 40-heptad (40H) class predominated across Vibrionales, whereas shorter variants such as 28H were exceedingly rare (only four instances), indicating strong selective constraints on receptor length and array assembly (Figure 6A, 6D). Importantly, no significant correlation was observed between MCP abundance and genome size (r=0.4, p= 6.93×10^-6^), supporting the notion that MCP expansion is not a by-product of genome scaling but likely reflects adaptive diversification driven by ecological complexity and niche specialization (Figure 6B), which is further supported by the disproportionately high MCP counts observed in members of the Gazogenes and Cholerae clades.

**Figure 6:**
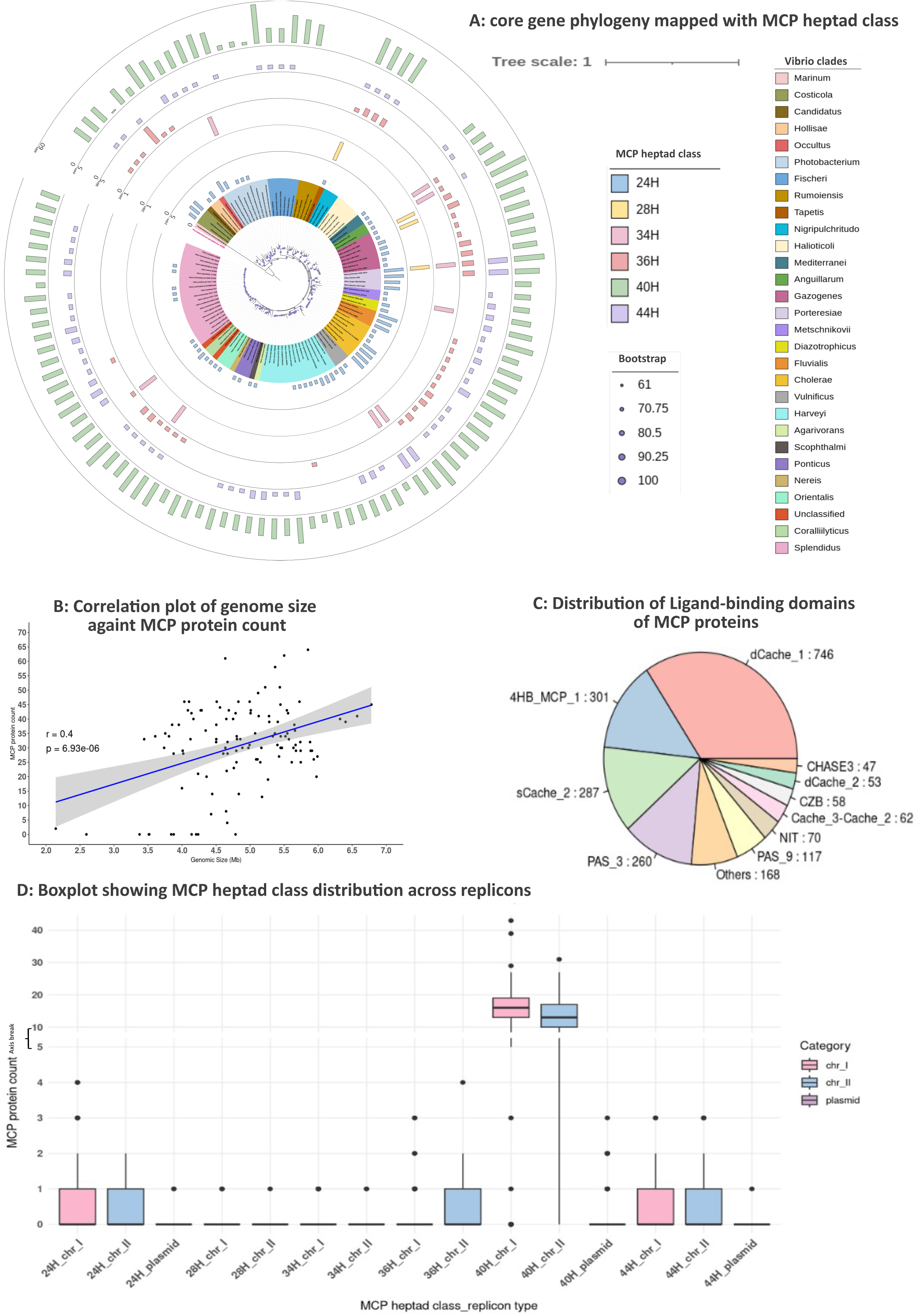
Distribution and structural diversity of methyl-accepting chemotaxis proteins (MCPs) across Vibrionales isolates. **(A)** Heptad class composition of MCP proteins mapped onto the core-genome phylogeny; each heptad class is represented by a distinct colour, with bar lengths indicating MCP counts per isolate. **(B)** Correlation between genome size and MCP protein count, calculated using cor.text function in R. **(C)** Pie chart showing the distribution of ligand-binding domain (LBD) types among MCP proteins. **(D)** Box plots depicting the distribution of MCP heptad classes across replicon types; the 40-heptad (40H) class is the most abundant across both chromosomal replicons.

Domain architecture analysis revealed extensive diversity in periplasmic ligand-binding domains (LBDs), with twenty-four distinct LBD families identified across Vibrionales MCPs (Figure 6C). Among these, dCache_1 (746, 33%), 4HB_MCP_1 (13%), sCache_2 (12%), and PAS_3 (11%) were the most abundant, collectively accounting for 69% of sensory inputs. The distribution of these major sensory domains across the major clades (species count more than 5) revealed that dCache_1 domain, known to detect a wide spectrum of ligands, proteinogenic amino acids, polyamines, quaternary amines, purines, organic acids (OAs), sugars, quorum sensing signals, or inorganic ions (Feng et al., 2022), is predominately present across these clades (Figure S7). The presence of MCP proteins on different replicons further highlights their evolutionary plasticity. We also report twenty-four plasmid-encoded MCPs, which represent a dynamic reservoir for horizontal gene transfer, enabling rapid acquisition of novel sensory capabilities. Such modular expansion of sensory repertoires is consistent with the “plug-and-play” model of chemosensory evolution (Lindemann et al., 2016), wherein MCPs can be independently gained or lost without disrupting core signaling machinery. The observed diversity of MCP repertoires reflects the broad ecological distribution of Vibrionales and their capacity to respond to varied environmental stimuli.

We further investigated the diversity within a single strain, *V. cholerae* O1 biovar EI Tor N16961, to observe their ligand binding patterns and sequence homology. The unrooted MCP protein phylogeny resolved into distinct clades corresponding to LBD diversity (Figure S6), showcasing that different LBD functions might have evolved at different times via duplication and later got modified to sense similar or different types of molecules.

## Discussion

Chemosensory systems represent one of the most extensively characterized signal transduction paradigms in bacteria. However, their detailed mechanistic and evolutionary insights have largely been derived from a limited number of model organisms, including *Escherichia coli* (Baker et al., 2006) and *Thermotoga maritima* (Perez & Stock, 2007), where canonical architectures and signaling principles have been established. In contrast, despite their ecological and clinical importance, CSS organization and functional diversification in broad taxonomy groups such as Vibrionales, particularly across non-model species, remain comparatively underexplored. Previous studies in *V. cholerae* have identified three major chemosensory clusters (F6, F7, and F9), with the F6 system being the best characterized due to its direct role in flagellar motility and host colonization (Ortega et al., 2020). Several chemotactic genes from F6 (cluster II), such as *mlp24*, *cheZ*, *cheA,* and *cheY,* are essential for the full induction of cholera toxin or the positive transcriptional regulator ToxT upon mouse infection (Hiremath et al., 2015). F9 (cluster I) proteins are known to be expressed only under particular conditions that bacteria might encounter upon infection (Hiremath et al., 2015). It has been hypothesized that F9 (cluster I) plays a role in sensing and signaling within the host intestine (anaerobic environment), potentially regulating virulence in that anaerobic host environment (Hiremath et al., 2015). The proteins of F9 and F6 were reported to be expressed during the Viable but Non-culturable (VBNC) state of *V. cholerae* (Brenzinger et al., 2019). However, the overall functional relevance of F7 and F9 systems also remains incompletely resolved.

This study presents the most comprehensive comparative genomic framework of chemosensory systems in Vibrionales to date, integrating 116 curated representative genomes along with a large-scale RefSeq and MAG dataset of ∼10,000 organisms. Our analyses resolve four discrete CSS lineages, F6, F7, F8, and F9, that constitute a conserved signaling toolkit across the whole order. The consistent identification of these four CSS types across independent, phylogenetically broad datasets of ∼10,000 genomes and MAGs argues strongly against sampling bias and supports the interpretation that Vibrionales operate with an evolutionarily constrained core set of chemosensory architectures. Critically, this study identifies the F8 system, which has not been reported earlier in Vibrionales experimental studies, showcasing the limits of organism-centric functional studies (*V. cholerae* in this case) and the power of large-scale comparative genomics to reveal cryptic regulatory modules.

F8 CSS has been extensively described in phylogenetically distant bacteria, including *Rhodobacter sphaeroides* (phylum Pseudomonadota) *and Borrelia burgdorferi* (phylum Spirochaetota), where its functions span motility regulation to poorly defined processes (Gumerov et al., 2021). Our CheA homolog phylogeny revealed that F8-CheA homologs are predominantly present in phylum Pseudomonadota (especially in class Gammaproteobacteria) and sparsely distributed across Betaproteobacteria, Desulfovibrionia, Desulfuromonadia, and Spirochaetia classes. Overall, the evolutionary history of F8 in Vibrionales can be putatively explained by ancestral retention followed by lineage-specific loss, rather than by recent horizontal acquisition. However, additional and deeper phylogenetic analyses are required to distinguish between these evolutionary scenarios.

However, F9 evolutionary trajectory exhibits putative signatures of horizontal gene transfer via its nearby clades and their modular architecture. The selective retention of F8 and F9 CSSs in a few organisms within specific clades raises the hypothesis that it mediates niche-adapted sensory functions, potentially linked to host association, nutrient gradients, or abiotic stress, which confer fitness benefits in particular ecological contexts. Functional dissection through targeted deletion of F8 and F9 CSS-associated components will be necessary to test this model directly.

The replicon-level organization of CSS clusters reveals a functional hierarchy that mirrors the biology of multipartite Vibrionales genomes. The predominant localization of F6 on chromosome I is consistent with its established role in flagellar motility and host colonization. Such conserved presence is also indicative of strong purifying selection acting on a core fitness determinant. All F9 clusters are located on chromosome I; however, F7 and F8 are sporadically distributed across both chromosomes, and the occasional presence of CSS components on plasmids reflects greater adaptive potential of organisms harbouring these systems. The plasmid-encoded F6-like cluster identified in *Photobacterium toruni*, which notably lacks CheB, represents a particularly intriguing case of possible functional repurposing, and may reflect an intermediate stage in the evolutionary decay of an accessory signaling module. Similarly, clade-specific arrangements, such as the coexistence of F7 and F8 in *V. gallaecicus* and the exclusive retention of F8 in *V. porteresiae*, illustrate how gene gain, loss, and selective retention fine-tune CSS composition at the level of individual species, likely in response to micro-niche ecological pressures.

At the protein level, CheA proteins across all four lineages retain the canonical histidine kinase domain organization; however, insertions between the P2 and P3 domains in F8 and F9 CheA proteins indicate structural and functional divergence. These insertions may alter interaction surfaces with CheW scaffold proteins, MCPs, or downstream response regulators, with consequences for signaling kinetics or pathway specificity. Domain-level insertions of this kind might act for functional innovation within conserved modular signaling scaffolds, and their presence in F8 and F9 warrants structural characterization to resolve their mechanistic significance. It remains to be explored whether the extra inserted region between P2 and P3 of CheA-F9 is functionally linked to the MCPs associated with the F9 cluster. Given that chemosensory array formation requires coordinated interactions between CheA and MCP proteins, this structural feature may play a role in array assembly. Notably, the specialized chemoreceptor DosM has been reported to be essential for the formation of cytoplasmic F9 arrays (Briegel et al., 2016), suggesting a potential structural and functional link between this structured region of CheA-F9 and MCP proteins, which needs further experimental validation. It was reported that the F9 class is rarely present alone and is most often encoded in a genome that also has other F classes (Mo et al., 2022). Such co-occurrence suggests that F9 may function in coordination with other chemosensory pathways rather than operating independently. Such an arrangement could reflect functional specialization, where F9 complements other CSS clusters by sensing distinct environmental cues.

Extensive expansion of MCP repertoires far beyond the number of core signaling modules is a significant feature of Vibrionales CSS organisation, which is well known across other organisms, i.e., Myxobacteria, *Pseudomonas*, etc (Ortega et al., 2017; Sharma et al., 2018). MCPs are predominantly genomically dispersed, rather than co-localized with CSS gene clusters, and are largely absent from F6 loci, which supports a model of functional decoupling between sensory input and downstream signaling. Following this, multiple MCPs from diverse genomic contexts can feed into shared signal transduction arrays, enhancing integration capacity and enabling combinatorial sensing of complex environmental signals. Computational predictions in *P*. *aeruginosa* PAO1 have previously revealed that 23 diverse chemoreceptors feed into a single Che I/F6 pathway, controlling chemotaxis (Ortega et al., 2017). The elevated MCP abundance in these ecologically diverse clades suggests that sensory repertoire expansion is driven by environmental complexity rather than genome size, consistent with patterns reported across chemosensory-rich proteobacteria (Ortega et al., 2017). Experimentally characterized examples reinforce this view: VfcA in *Aliivibrio fischeri* mediates serine amino acid sensing (Brennan et al., 2013; Everiss et al., 1994), VfcB and VfcB2 detect short-and medium-chain aliphatic (or fatty) acids (Nikolakakis et al., 2016), and MCP-like proteins including AcfB and TcpI in *V. cholerae* contribute directly to intestinal colonization, establishing a mechanistic link between chemoreceptor diversity, metabolic sensing, and pathogenic potential (Chaparro et al., 2010). *V. cholerae* MCP protein Mlp45 (VcAer2), associated with F7 (cluster III), senses the oxygen concentration (Greer-Phillips et al., 2018). *V. cholerae* utilizes a dedicated MCP protein MCP^DRK^ (VC1313) to sense D-amino acids as a repellent signal via signalling to CSS cluster II/F6 to control motility (Irazoki et al., 2023).

Altogether, our findings reveal four types of Vibrionales chemosensory networks, which are organized as a two-tier architecture: 1) a conserved signaling core of F6-type CSS, which has been subjected to an ongoing purifying selection, and 2) a dynamically evolving permutation-combination system of F7, F8, F9 accessory CSSs that expand based on the adaptive range of the sensory network. This organizational pattern is not unique to Vibrionales, as analogous tiered architectures have been described in *Pseudomonas aeruginosa*, which encodes a conserved Che1 core system alongside multiple accessory systems (Che2-Che5) linked to biofilm formation and virulence (Ortega et al., 2017), and in *Myxococcus xanthus*, where spatially segregated 8-12 chemosensory clusters mediate distinct behavioural outputs helping in its multicellular social behaviour (Sharma et al., 2018). However, what distinguishes Vibrionales is the tight coupling of this modularity to multipartite genome organization: the preferentially conserved localization of F6 on the primary chromosome, combined with the replicon-level plasticity of accessory CSSs and their occasional plasmid carriage, suggesting that genome architecture itself serves as an evolutionary substrate for chemosensory innovation. From a broader evolutionary perspective, this two-tier design mirrors principles observed in other modular prokaryotic signaling networks, such as two-component systems and second messenger cascades, where a conserved transduction core is repeatedly co-opted and diversified through the acquisition of peripheral input modules. The decoupling of receptor localization from core signaling components, and the plasticity in replicon-level gene distribution, enables rapid niche adaptation without compromising the fidelity of core chemotactic output. Resolving the specific contributions of F8 and characterizing the extra inserted region within F8/F9 CheA proteins will be essential next steps toward a mechanistic understanding of CSS diversification and its role in environmental adaptation and pathogenesis across Vibrionales.

## Supplementary figure legends

**Figure S1: (A)** Correlation plot showing the association between genome size of large chromosome and small chromosome. **(B)** Correlation plot depicting the association between CDS count on respective replicon and genome size of respective replicon. **C and D** represent the correlation plots between total ribosomal RNA count and genome size and total tRNA count and genome size, respectively.

**Figure S2:** Panels showing the correlation plots between orphan histidine kinase, orphan response regulator, and hybrid TCS and chromosome genome size. **A, B**, and **C** represent the correlation plots between orphan histidine kinase and chromosome size, orphan response regulator and chromosome size, and hybrid TCS and chromosome size, respectively.

**Figure S3:** Boxplots A and B depicting the distribution of CheA and MCP across *Vibrio* clades, respectively. The numbers in brackets represent the number of species in each clade.

**Figure S4:** Core gene phylogeny representing the distribution of each CSS cluster across different replicons in Vibrionales.

**Figure S5:**
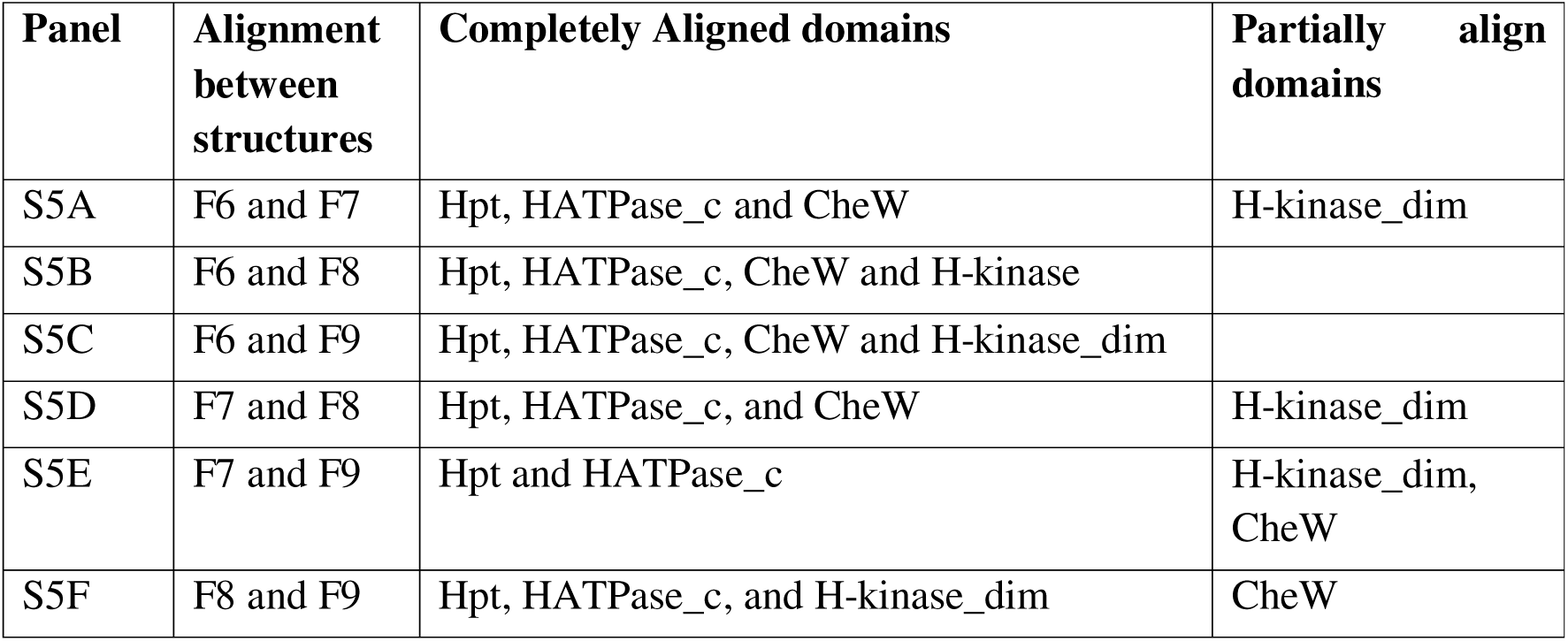
Panels showing the structural alignment for all four CheA proteins. F6, F7, F8, and F9 are represented in pink, yellow, orange, and cyan, respectively. The TM-align score for each structural alignment is shown in the table.

**Figure S6:** The unrooted MCP protein phylogeny constructed using forty-six identified MCP proteins of *V. cholerae* O1 biovar EI Tor N16961 strain in this study. The heptad class and domain architecture for each MCP protein are mapped on the phylogeny using iTOL v6. The sequences were aligned using MUSCLE v5.1, and the alignment block length of 1,670 was used for this LG+F+R5 model-based ML phylogeny and visualized in iTOL.

**Figure S7:** The boxplot showing the distribution of four abundantly present ligand-binding domains of MCP protein across the top 6 *Vibrio* clades (clades having species count more than 5).

## Supplementary table legends

**Supplementary table S1:** Table showing the ecological habitat, nature of association of 116 Vibrionales isolates along with relevant reference.

**Supplementary table S2:** Table showing the distribution of GC content, rRNA, tRNA, and protein-coding gene count across different replicons among Vibrionales isolates along with their NCBI accession number, clade name, and genus.

**Supplementary table S3:** Table representing the distribution of all chemosensory system proteins across different replicons among Vibrionales isolates with their NCBI accession number, clade name, and genus.

**Supplementary table S4:** Table showing the count of each CSS cluster along with CSS proteins among Vibrionales members, along with their NCBI accession number, clade name, and genus.

**Supplementary table S5:** Table showing the different CSS cluster categories across Vibrionales members.

**Supplementary table S6:** Table showing the total, minimum, maximum, average, and median MCP protein count across each *Vibrio* clade

## Data availability statement

Publicly available datasets were analysed in this study. The NCBI accession numbers of these datasets are available in the “B” column in the supplementary table.

## Conflict of Interest

The authors declare that the research was conducted in the absence of any commercial or financial relationships that could be construed as a potential conflict of interest.

## Author contributions

SR: Investigation, Methodology, Resources, Validation, Data curation, Formal analysis, Visualization, Writing - original draft

GS: Conceptualization, Funding acquisition, Project administration, Supervision, Methodology, Resources, Validation, Writing - review & editing

## Funding

SR acknowledges the fellowship by the DST-INSPIRE Program of the Department of Science and Technology. GS is supported by the Start-up Research Grant (SRG) from the Anusandhan National Research Foundation (ANRF), India.

## Acknowledgments

The support and computational resources provided by PARAM Seva Facility under the National Supercomputing Mission (NSM), Government of India at the Indian Institute of Technology Hyderabad are gratefully acknowledged. The authors would like to thank Jaimin Chodvadiya for providing a computational script that contributed to portions of the data analysis performed in this study and Normada Hatimuria and Shashank Singh for performing the literature review of the ecological habitat of Vibrionales.

## Notes

### Competing Interest Statement

The authors have declared no competing interest.

### Summary of Updates

This version of the manuscript has been revised to substantially improve the Introduction, clarify the objectives and novelty of the study, and provide a broader overview of bacterial chemosensory systems and their evolution. The discussion of Vibrionales biology, quorum sensing, chemotaxis, and chemosensory signaling has been streamlined and updated with additional recent references. The classification, distribution, and evolutionary relationships of F6, F7, F8, and F9 chemosensory systems have been further clarified, including their broader context within Gammaproteobacteria and the possible horizontal acquisition of F9 from Alphaproteobacteria. Figures and phylogenetic trees have been revised for improved readability, with bootstrap support added to phylogenetic analyses. Structural analyses have also been strengthened by comparison of predicted CheA structures with the experimentally characterized Escherichia coli CheA using TM-align. Methods and relevant interpretations have been revised accordingly, and additional references have been incorporated throughout the manuscript.

