## Supplementary Figures for "Chemosensory diversity across the order Vibrionales reveals a conserved core and three accessory signalling systems"

**S1**

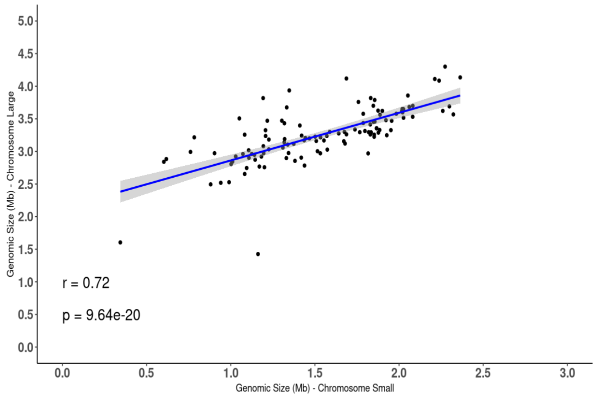


**A: Correlation plot of genomic sizes of large and small chromosome**

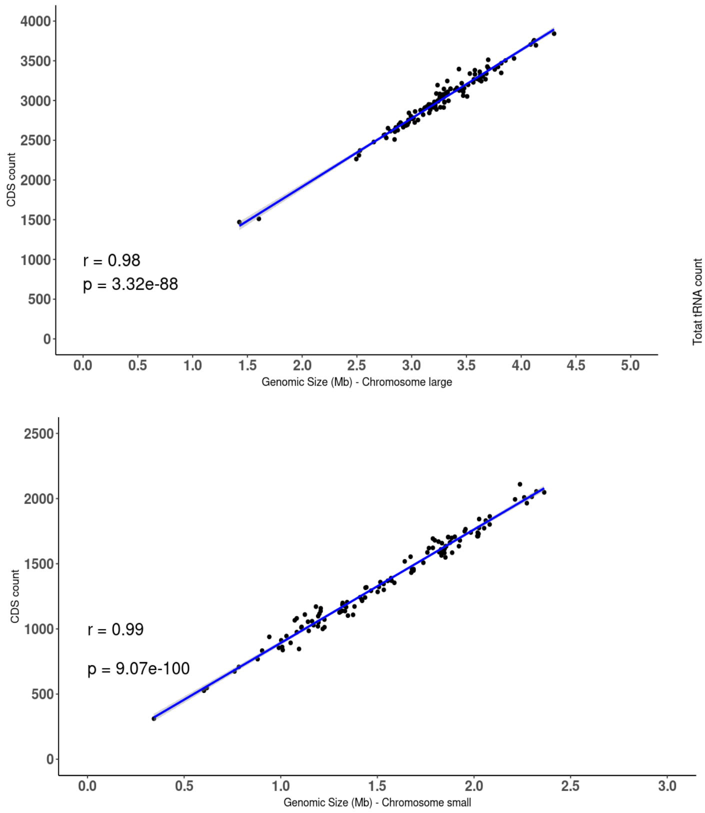


**B: Correlation plots (CDS count against genome size) for large and small chromosome**

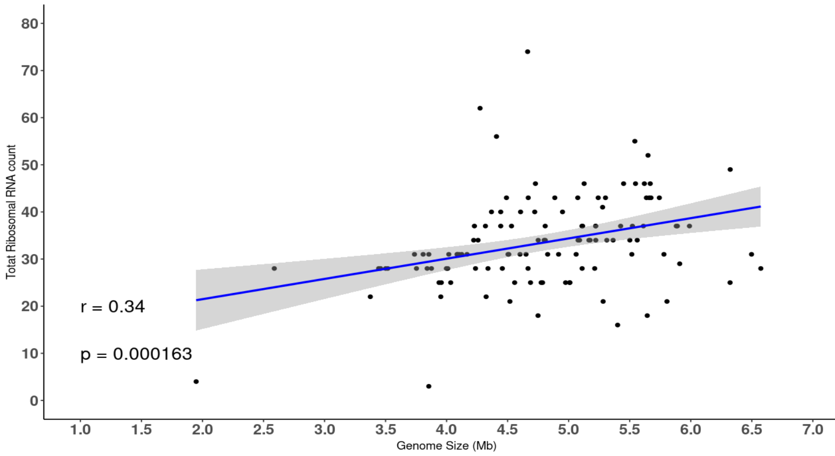


**C: Correlation between total ribosomal RNA count and genome size**

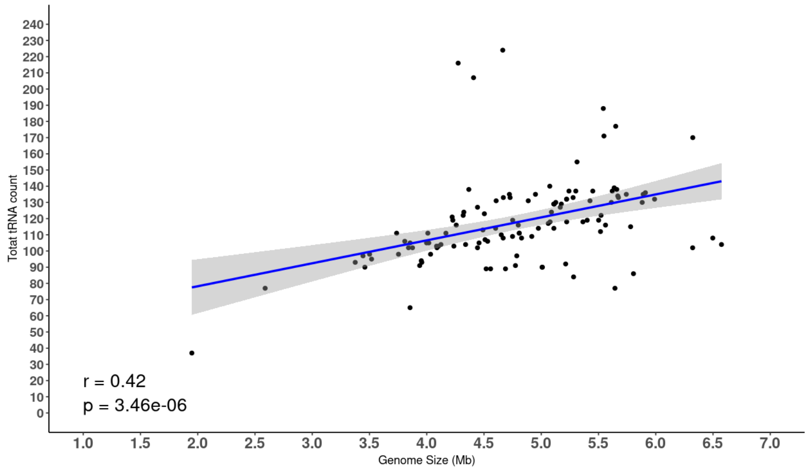


**D: Correlation between total tRNA count and genome size**

### Multipartite genome and extensive RNA gene repertoire reflect niche adaptation in Vibrionales:

Apart from diverse ecological habitats, the presence of a multi-partite genome is a peculiar characteristic of Vibrionales members. All these organisms have two chromosomes with a few optional plasmids which encompasses significant differences between their genomic sizes. Chromosome I is typically 2-3 times larger in genomic size (average 3.25 Mb) as compared to chromosome II (average of 1.53 Mb) (Figure 1D). Irrespective of different genomic sizes, the GC content for both chromosomes is almost the same, i.e., ~45% (Figure 1E). The total genome size of Vibrionales members ranges from 4 to 6 Mb (average and median of 4.8 Mb), aligning with the upper spectrum in the entire bacterial kingdom (average of 3.35 Mb, median of 2.9 Mb, for 7,29,560 NCBI genomes (2024 dataset)). Notably, *Candidatus Enterovibrio luxaltus* CC26 represents an extreme case of genome reduction within the order, with a total genome size of 2.14 Mb, partitioned into a disproportionately small chromosome II (~343 kb), indicative of potential genome streamlining associated with niche specialization or host dependency.

A significant positive correlation between the sizes of chromosome I and chromosome II ($r = 0.72$) indicate coordinated genome expansion across replicons, suggesting that genome growth in Vibrionales is not restricted to the primary chromosome but involves parallel scaling of secondary replicons (Figure S1A). We further report a strong positive correlation between genomic sizes of both chromosomes and their respective total protein coding genes, as indicated by high value of correlation coefficient (r=.0.99 and r=0.98), and low p-value (Figure S1B), indicating that genome expansion is predominantly driven by proportional increases in protein-coding gene content rather than accumulation of non-coding DNA. Overall, the number of coding genes of chromosome I (average is 2993) is larger than chromosome II (average is 1359), supporting a functional partitioning model in which core cellular processes are enriched on chromosome I, while chromosome II likely harbors accessory and niche-adaptive genes. Within Vibrionales order, coding capacity (CDS counts) varies widely, ranging from 2,019 in *Candidatus Enterovibrio luxaltus* CC26 to 6,028 in *V. penaeicida* IFO 15640T, reflecting substantial genomic diversity within the order (Table S2).

Analysis of RNA gene distribution revealed considerable variability in ribosomal RNA (rRNA) and transfer RNA (tRNA) across Vibrionales (Table S2). While the average rRNA copy number is ~33, nine members of the *Photobacterium* clade exhibit exceptionally elevated counts (mean ~182), suggesting lineage-specific amplification potentially linked to rapid growth rates or fluctuating environmental conditions requiring enhanced translational capacity. Similarly, tRNA gene counts are also consistently high (average ~117), supporting vigorous translational machinery across diverse ecological habitats (Table S2). It must be noted that neither rRNA nor tRNA counts shows a significant correlation with genome size ($p \approx 0.34$ and 0.42, respectively) (Figure S1C, Figure S1D), indicating that their RNA-level expansion might be regulated by functional/physiological requirements rather than in accordance with the genome size. Interestingly, while most Vibrionales plasmids lack rRNA operons, selected species such as *Candidatus Enterovibrio luxaltus* CC26, *Photobacterium toruni* WD2103, and *V. scophthalmi* VS-05 encode tRNA genes on plasmids, suggesting their potential roles in horizontal gene transfer, or adaptive responses to environmental stress, further making them significant evolutionary drivers.

S2

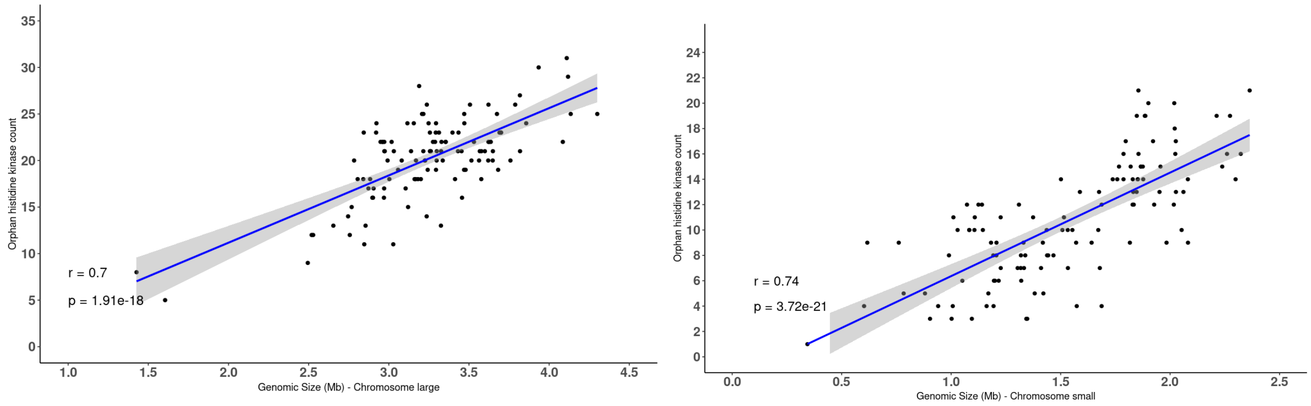


**A: Correlation between orphan histidine kinase and chromosome genome size**

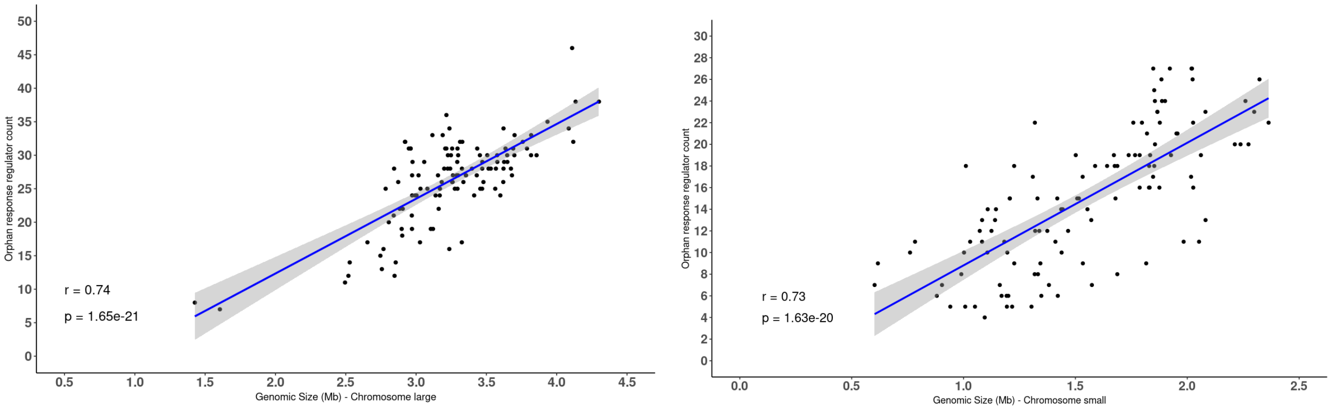


**B: Correlation between orphan response regulator and chromosome genome size**

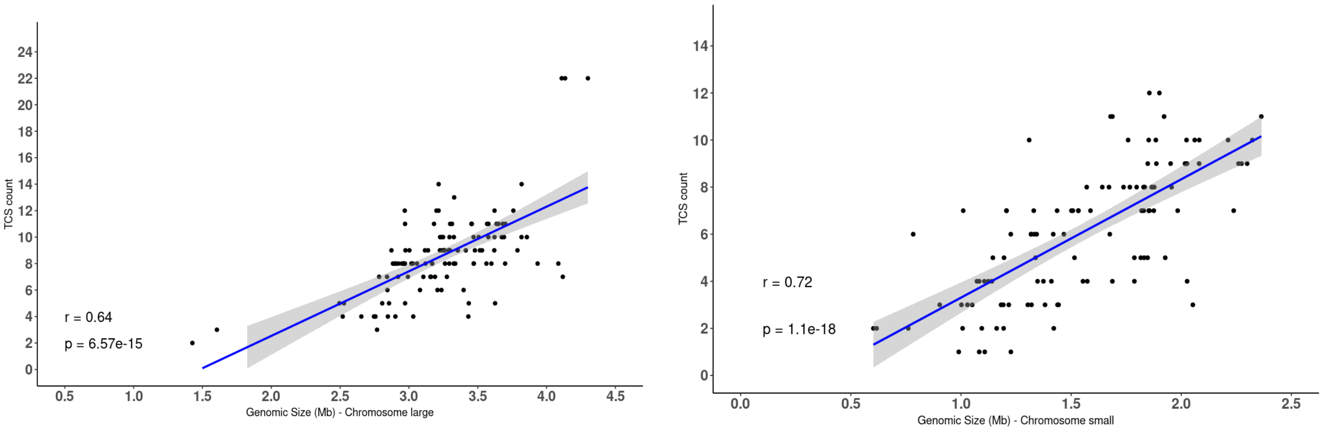


**C: Correlation between Hybrid Two-component system (TCS) and chromosome genome size**

**S3**

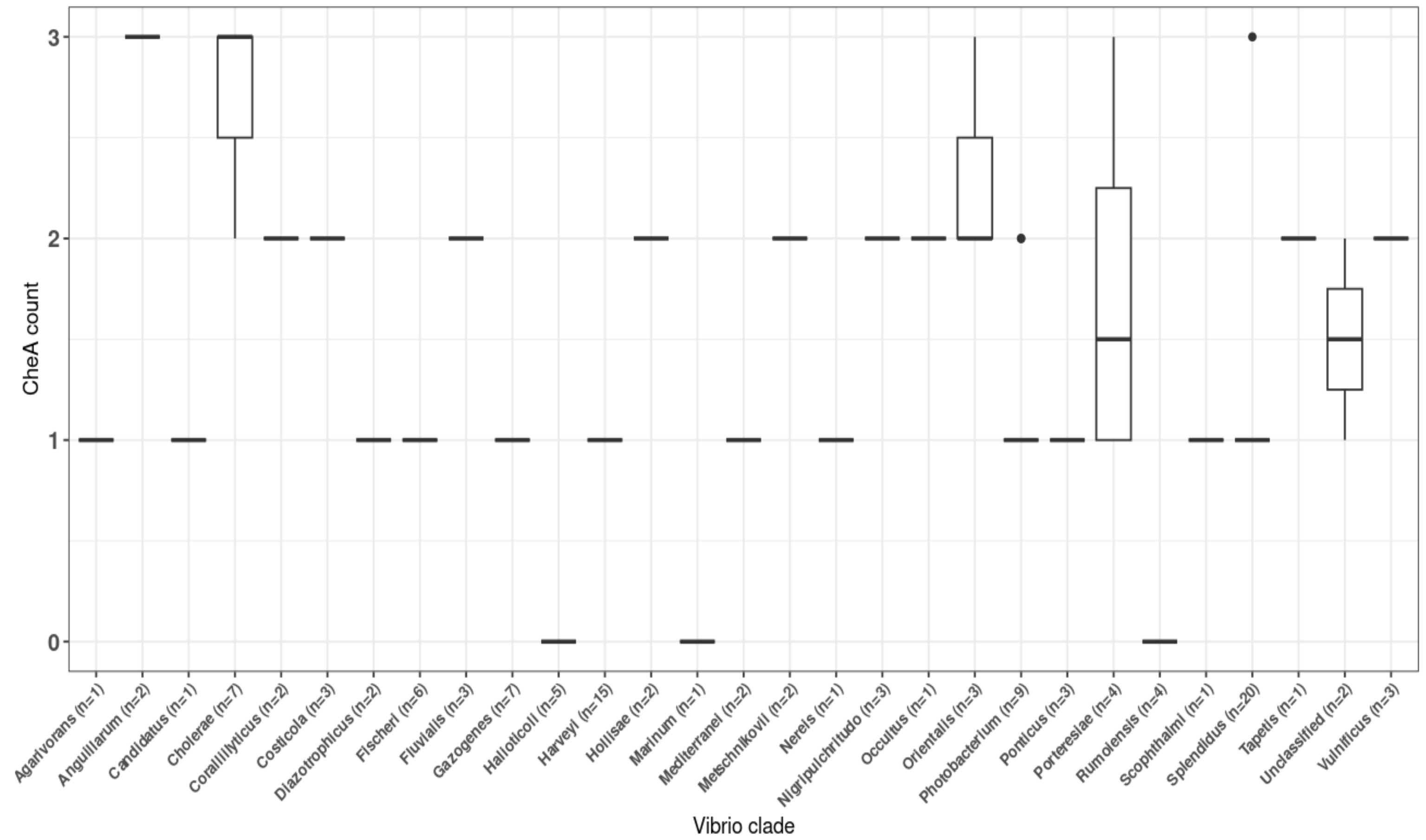


**A: Distribution of CheA proteins acorss Vibrio clades**

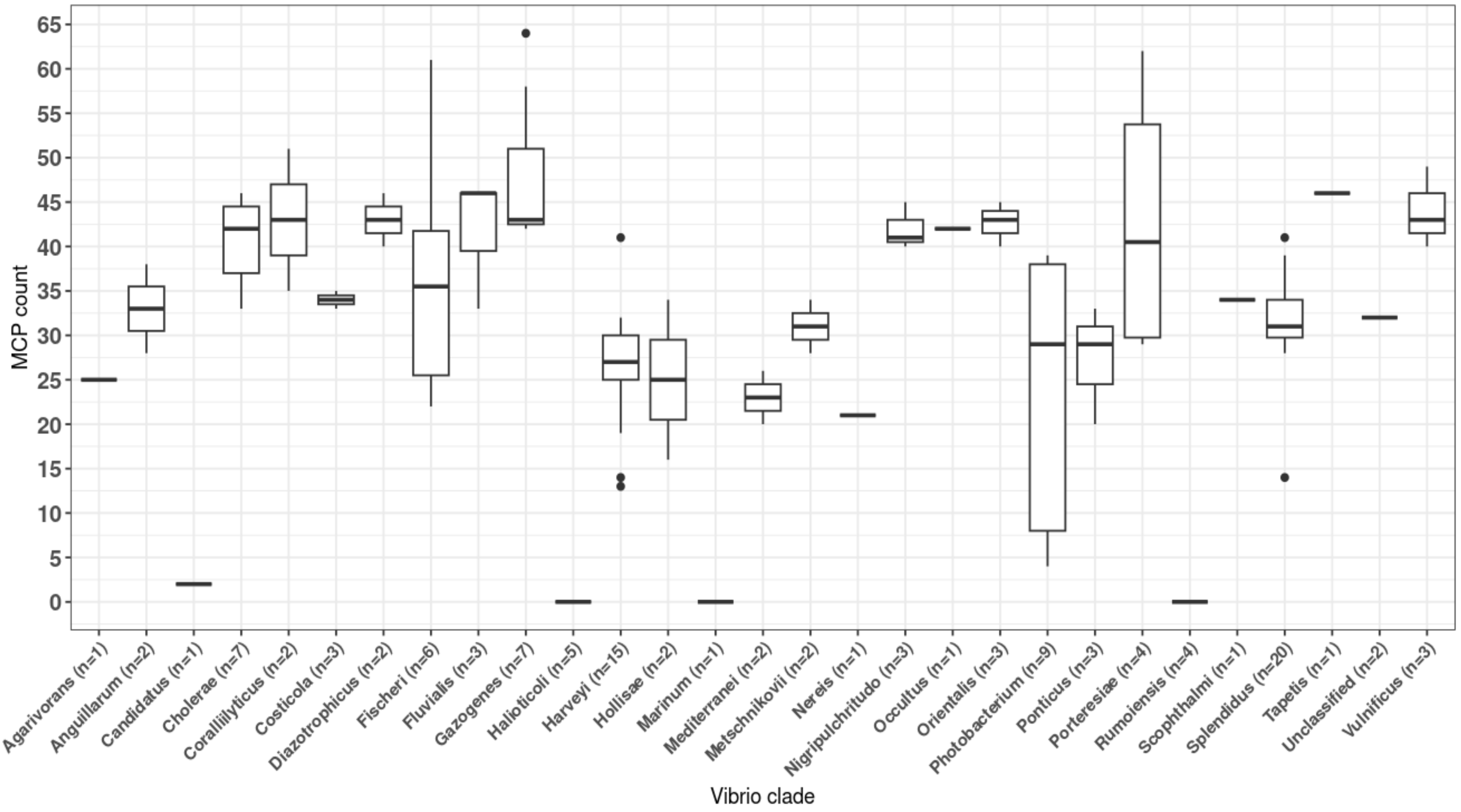


**B: Distribution of MCP proteins acorss Vibrio clades**

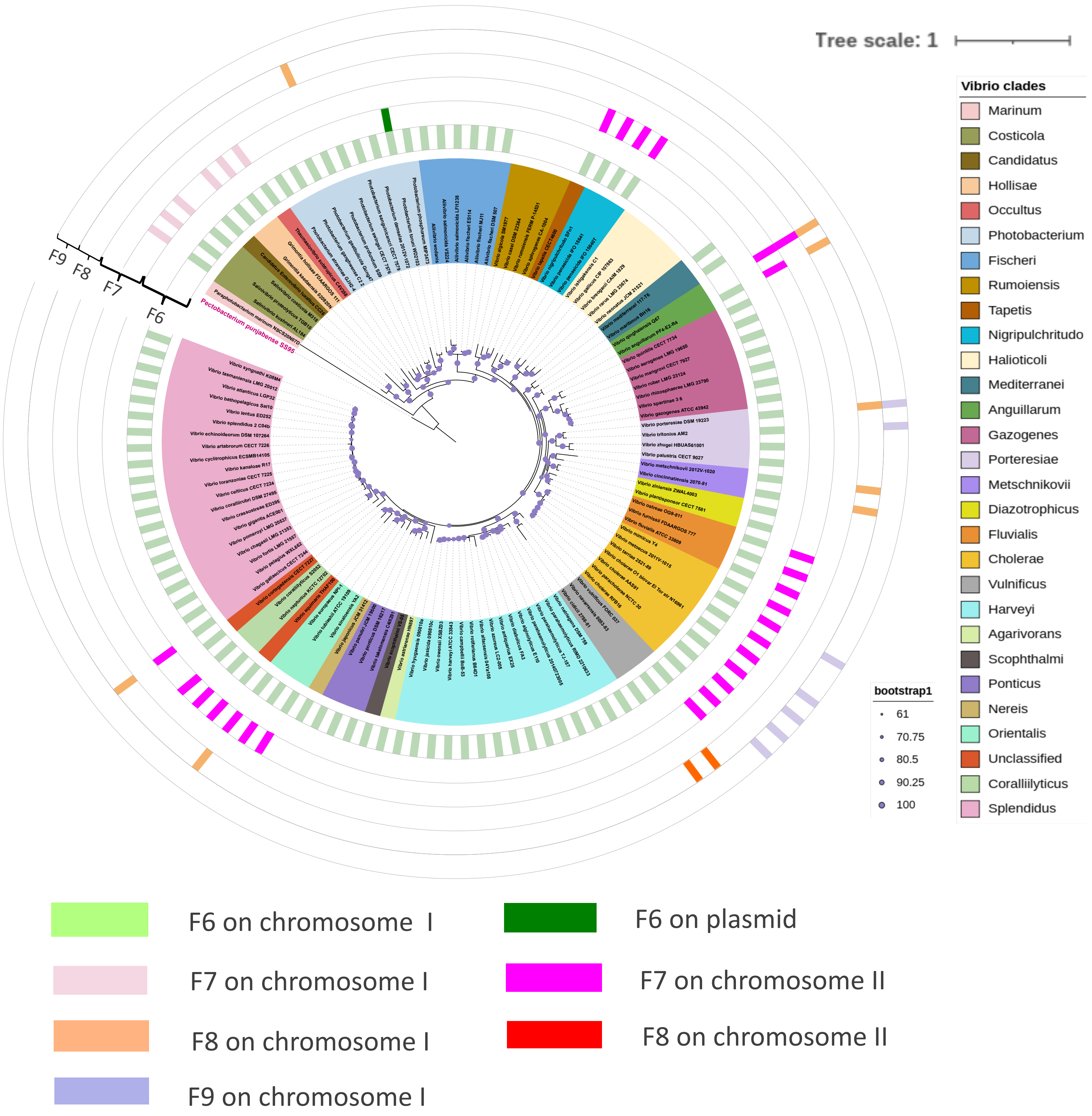


**S4: Core gene phylogeny representating the distribution of each CSS cluster across different replicons in Vibrionales**

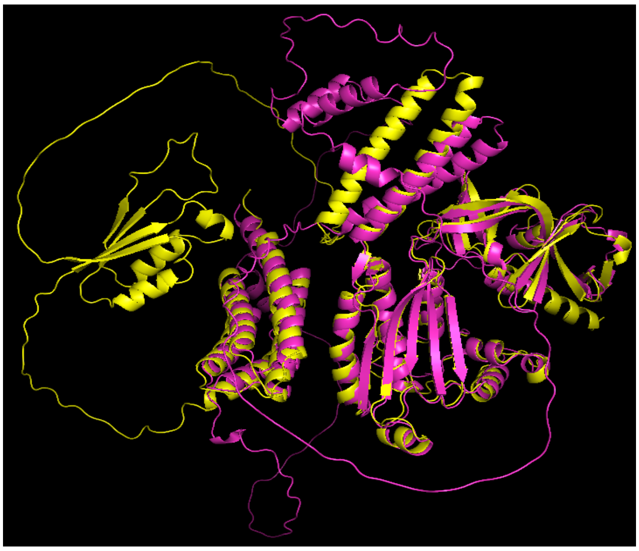

S5A: Yellow: F7 Pink: F6

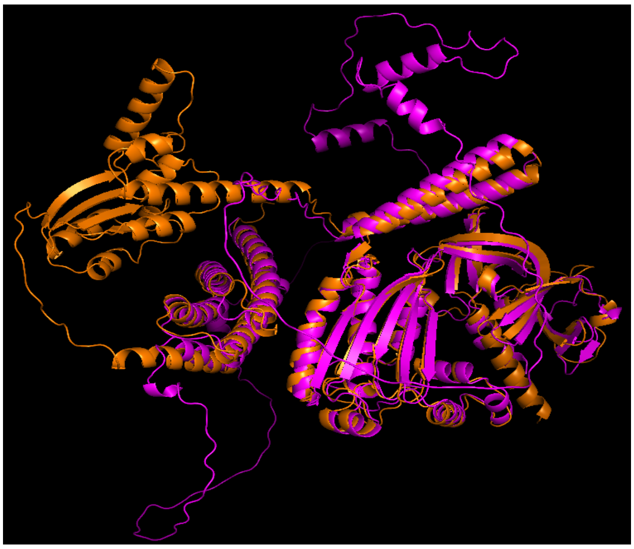

S5B: Pink: F6 Orange: F8

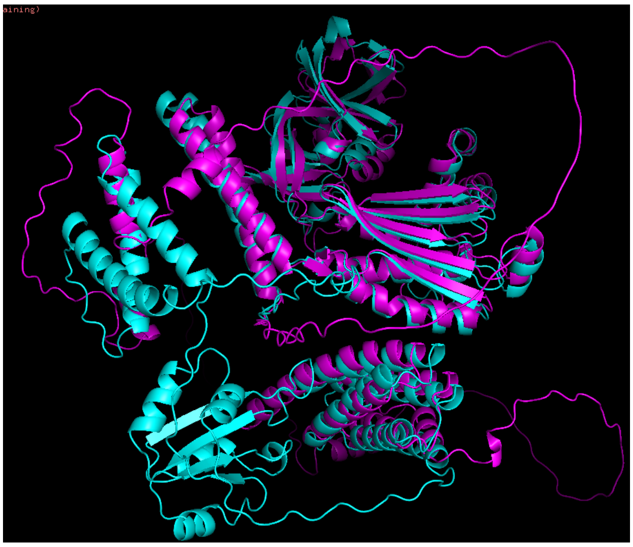

S5C: Pink: F6 Cyan:F9

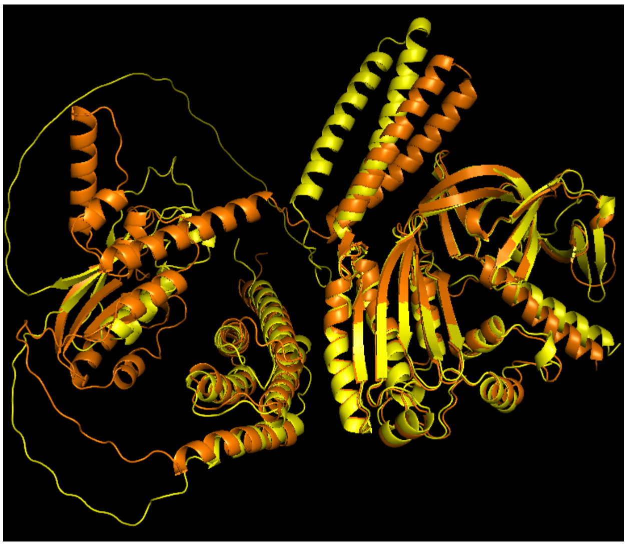

S5D: Yellow:F7 Orange:F8

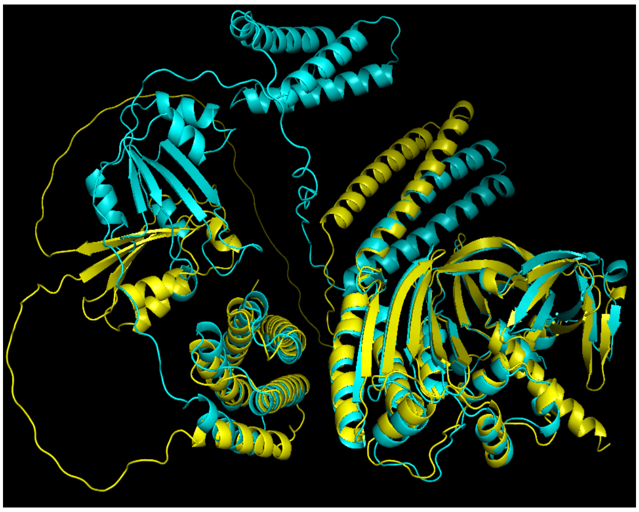

S5E: Yellow:F7 Cyan:F9

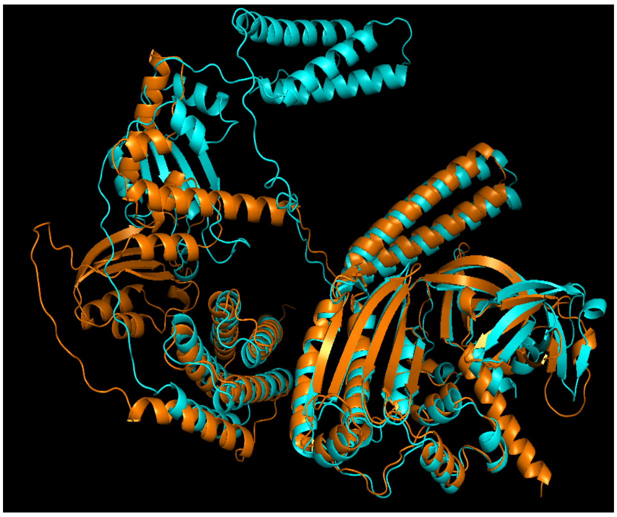

S5F: Orange:F8 Cyan:F9

**S5: Structure alignment of CheA proteins along with the RMSD value for each alignment**

| Sr.No. | Structural alignment of CheA proteins | TM-score (normalized by length of chain 1) | TM-score (normalized by length of chain 2) |
|---|---|---|---|
| 1. | F6 and F7 | 0.63 | 0.66 |
| 2. | F6 and F8 | 0.67 | 0.67 |
| 3. | F6 and F9 | 0.65 | 0.63 |
| 4. | F7 and F8 | 0.75 | 0.71 |
| 5. | F7 and F9 | 0.68 | 0.62 |
| 6. | F8 and F9 | 0.69 | 0.66 |

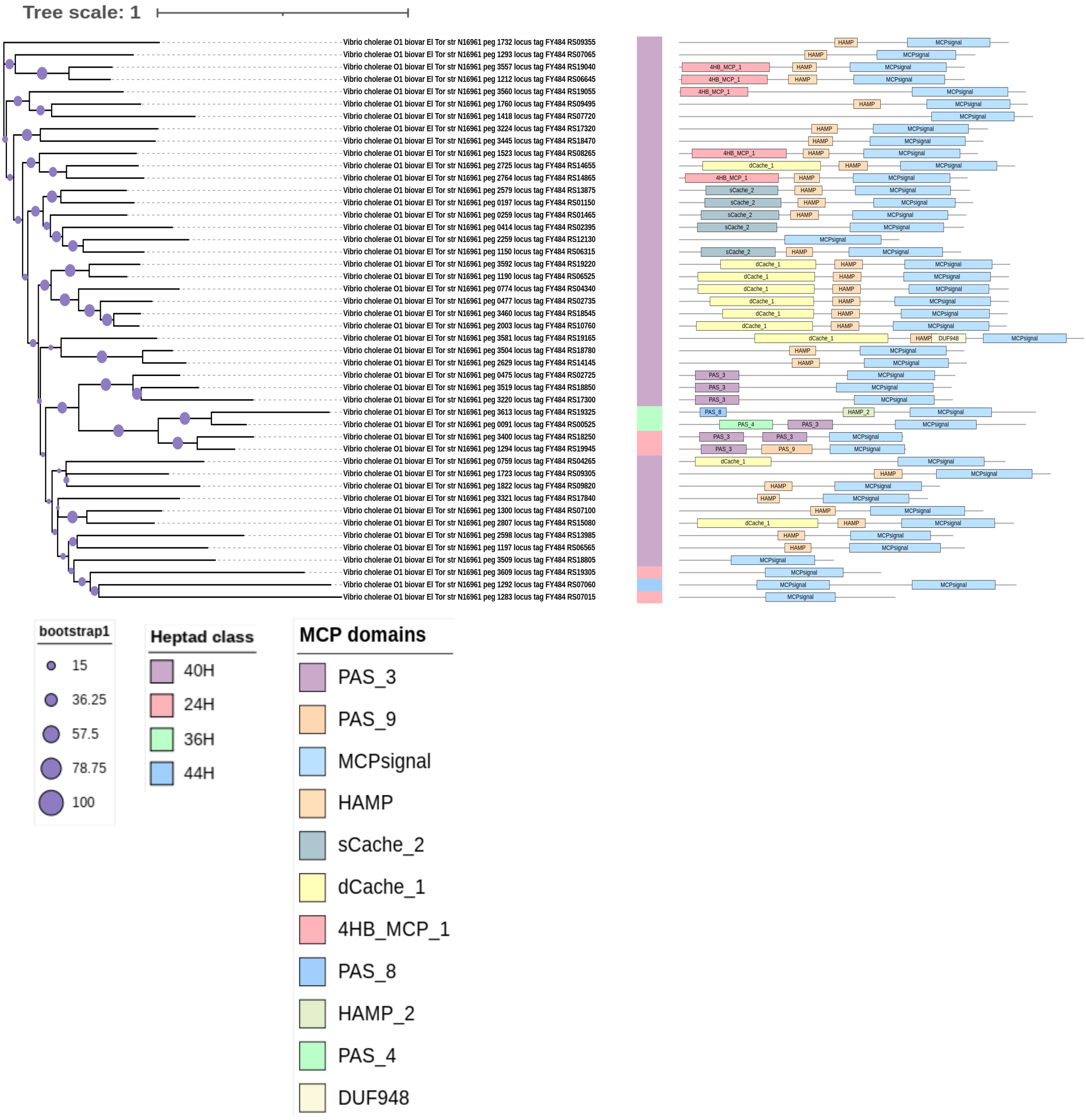


**S6: The Unrooted MCP protein phylogeny is constructed using the MCP proteins of *Vibrio cholerae* O1 biovar El Tor N16961 strain.**

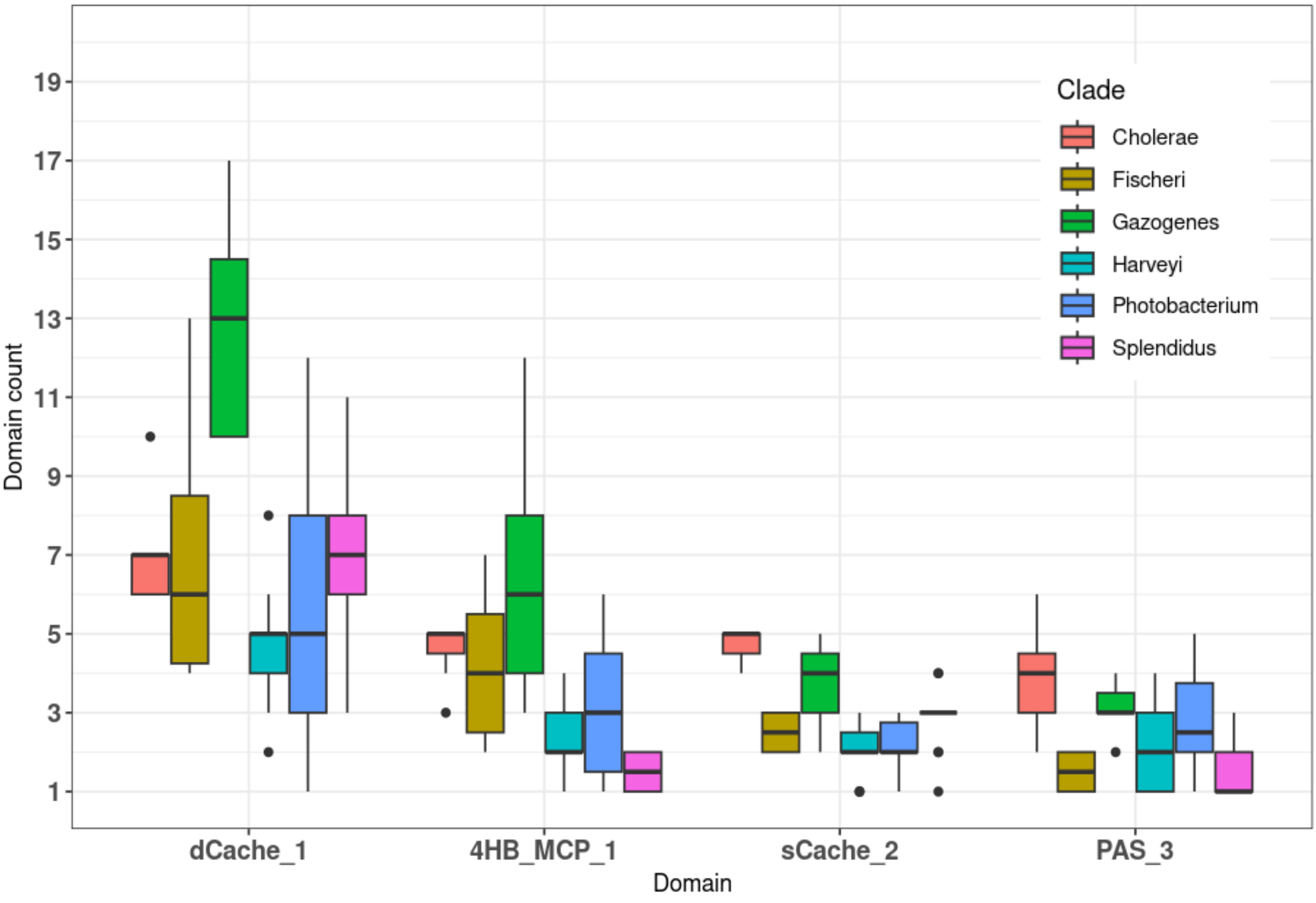


**S7: The boxplot showing the distribution of four ligand binding domains (X-axis) across top 6 *Vibrio* clades (clades having species count above 5)**
